# Mind the gap: performance metric evaluation in brain-age prediction

**DOI:** 10.1101/2021.05.16.444349

**Authors:** Ann-Marie G. de Lange, Melis Anatürk, Jaroslav Rokicki, Laura K.M. Han, Katja Franke, Dag Alnæs, Klaus P. Ebmeier, Bogdan Draganski, Tobias Kaufmann, Lars T. Westlye, Tim Hahn, James H. Cole

**Affiliations:** LREN, Centre for Research in Neurosciences- Dept. of Clinical Neurosciences, CHUV and University of Lausanne, Lausanne, Switzerland; NORMENT, Institute of Clinical Medicine, University of Oslo, & Division of Mental Health and Addiction, Oslo University Hospital, Oslo, Norway; Dept. of Psychiatry, University of Oxford, Oxford, UK; Centre for Medical Image Computing, Dept. of Computer Science, University College London, London, UK; Centre of Research and Education in Forensic Psychiatry, Oslo University Hospital, Oslo, Norway; Dept. of Psychiatry, Amsterdam University Medical Centers, Vrije Universiteit and GGZ inGeest, Amsterdam Neuroscience, Amsterdam, The Netherlands; Structural Brain Mapping Group, Dept. of Neurology, Jena University Hospital, Jena, Germany; Dept. of Neurology, Max Planck Institute for Human Cognitive and Brain Sciences, Leipzig, Germany; Tübingen Center for Mental Health, Dept. of Psychiatry and Psychotherapy, University of Tübingen, Tübingen, Germany; Dept. of Psychology, University of Oslo, Oslo, Norway; KG Jebsen Centre for Neurodevelopmental Disorders, University of Oslo, Oslo, Norway; Institute of Translational Psychiatry, University of Münster, Münster, Germany; Dementia Research Centre, Institute of Neurology, University College London, London, UK

**Keywords:** Brain-age prediction, Neuroimaging, Machine learning, Statistics

## Abstract

Estimating age based on neuroimaging-derived data has become a popular approach to developing markers for brain integrity and health. While a variety of machine-learning algorithms can provide accurate predictions of age based on brain characteristics, there is significant variation in model accuracy reported across studies. We predicted age based on neuroimaging data in two population-based datasets, and assessed the effects of age range, sample size, and age-bias correction on the model performance metrics *r, R*^2^, Root Mean Squared Error (RMSE), and Mean Absolute Error (MAE). The results showed that these metrics vary considerably depending on cohort age range; *r* and *R*^2^ values are lower when measured in samples with a narrower age range. RMSE and MAE are also lower in samples with a narrower age range due to smaller errors/brain age delta values when predictions are closer to the mean age of the group. Across subsets with different age ranges, performance metrics improve with increasing sample size. Performance metrics further vary depending on prediction variance as well as mean age difference between training and test sets, and age-bias corrected metrics indicate high accuracy - also for models showing poor initial performance. In conclusion, performance metrics used for evaluating age prediction models depend on cohort and study-specific data characteristics, and cannot be directly compared across different studies. Since age-bias corrected metrics in general indicate high accuracy, even for poorly performing models, inspection of uncorrected model results provides important information about underlying model attributes such as prediction variance.

## 1. Introduction

Brain-predicted age is increasingly used as a marker for structural brain integrity and health across normative and clinical populations [1, 2, 3, 4, 5, 6, 7, 8, 9, 10, 11, 12, 13, 14, 15, 16, 17]. Since brain structure is known to vary with age across the lifespan, machine learning (ML) regression models can be used to predict chronological age based on neuroimaging data [18, 19, 20, 21, 22]. Training a regression model on a wide range of magnetic resonance imaging (MRI) scans allows it to build a normative trajectory of brain differences across age, and condense a rich variety of brain characteristics into a single quantity per individual. Prediction models can then be applied to unseen data, providing an estimate of brain-predicted age for each individual in the dataset. The difference between an individual’s brain-predicted and chronological age (*brain age delta*) provides a proxy for deviations from expected age trajectories, and has been associated with clinical risk factors [10, 11, 23] as well as neurological and neuropsychiatric conditions [2, 9, 13, 20, 24, 25, 26, 27, 28, 29]. Brain age delta estimates have also been linked to biomedical variables and lifestyle factors in healthy population cohorts [3, 22, 11, 12, 30, 31, 32], and the overall evidence supports the use of brain-predicted age as a surrogate marker for brain integrity and health [21].

A number of recent studies show that ML algorithms can predict age based on MRI data with high accuracy e.g. [9, 33, 34, 26]. However, in addition to differences in feature sets included [10, 11, 35], training and test set characteristics such as size and age range [35, 10] can lead to considerable variation in model performance metrics across studies. Prediction accuracy is commonly evaluated using the correlation between brain-predicted and chronological age (*r*) or *R*^2^, in addition to Root Mean Squared Error (RMSE) and Mean Absolute Error (MAE). While these metrics are useful for comparing different algorithms applied to the same dataset, the comparison of model performance across studies is less straightforward. For example, the correlation coefficient is reduced when measured in restricted ranges of a variable [36, 37], while the model error metrics RMSE and MAE depend on the distribution of the predicted variable, and will thus vary between studies with different cohort age ranges. Statistical corrections of overestimated predictions in younger subjects and underestimated predictions in older subjects can also have a large effect on model performance metrics. This phenomenon, which is commonly referred to as age-bias in brain age studies [38, 39, 40, 41, 42], occurs due to general statistical features of a regression analysis [38] (see Section 2.6). Age-bias correction ensures that any associations with other variables of interests are not driven by the age-dependence of the predictions. However, model performance metrics calculated post-correction may not always provide a relevant or valid representation of the initial model performance. This is important since the validity of brain-predicted age estimates depends on aspects such as sufficient variance in predictions, which is contingent on how well the initial model performs.

With an increasing number of studies using brain age prediction, there is a pressing need to establish a general understanding of model performance metrics, and how and why they may vary across studies. In this work, we ran age prediction models based on T1-weighted imaging data in two population-based datasets, and assessed the effects of age range, sample size, and age-bias correction on metrics that are commonly used to evaluate model accuracy; *r, R*^2^, RMSE, and MAE.

## 2. Materials and Methods

### 2.1. Datasets and data availability

The data were derived from UK Biobank (UKB) and the Cambridge Centre for Ageing and Neuroscience dataset (Cam-CAN). Sample demographics are provided in Table 1. The two datasets were chosen due to large sample size (UKB) and wide age range (Cam-CAN). The data are available through established access procedures for UKB (https://www.ukbiobank.ac.uk/researchers) and Cam-CAN (http://https://www.cam-can.org/index.php?content=dataset). The code used for running the age prediction models is available at https://github.com/amdelange/brainage.

**Table 1:** Sample demographics.

|  |  | UKB | Cam-CAN |
| --- | --- | --- | --- |
| **N** |  | 41,285 | 622 |
| **Age** | Mean ± SD | 64.15 ± 7.54 | 54.17 ± 18.40 |
|  | Range [years] | 45 - 82 | 18 - 87 |
| **Sex** | % Men | 47.36 | 50.64 |
|  | % Women | 52.64 | 49.35 |

### 2.2. MRI data acquisition and processing

A detailed overview of the UKB data acquisition and protocols is provided in [43] and [44], and the processing pipeline is available in [9]. For Cam-CAN, study protocols are available in [45] and [46]. For each of the datasets, global and regional measures of cortical volume, area, and thickness in addition to subcortical volume were extracted based on the Desikan-Killiany atlas [47] and automatic subcortical segmentation in FreeSurfer (version 5.3) [48]. After excluding participants with low-quality MRI data (based on outlier detection for UKB as described in [49], and outlier detection plus manual inspection for Cam-CAN as described in [23]), data from 41,285 and 622 participants were included for UKB and Cam-CAN, respectively.

### 2.3. Brain-age prediction

To estimate global brain age, we used the *XGBoost* regression algorithm (XGB; https://github.com/dmlc/xgboost), which is based on gradient tree boosting. XGB has demonstrated high performance in previous machine learning competitions [50], and has been used in a number of recent brain age studies [9, 10, 12, 31, 49, 51, 52]. Model parameters were set to *number of estimators* = 180, *max depth* = 3, and *learning rate* = 0.1, as determined based on previous grid searches in the two datasets [51, 53]. Models were run for i) the full UKB and Cam-CAN samples using 10-fold cross validation, ii) UKB subsets with different age range and sample sizes (see section 2.5), and iii) UKB and Cam-CAN samples where fractions of the data were randomly shuffled (see section 2.6). For each iteration, the MRI input features were scaled using the robust scaler from the scikit-learn library [54], which removes the median and scales the data according to the quantile range. To test whether choice of algorithm influenced the results, we repeated the UKB analyses in sections 2.5 and 2.6 using Linear Support Vector Regression (SVR; https://scikit-learn.org/stable/modules/generated/sklearn.svm.LinearSVR.html) with loss=‘epsilon insensitive’ and C=1.5.

### 2.4. Model performance metrics

Model performance metrics included the correlation between brain-predicted and chronological age (Pearson’s *r*), *R*^2^, RMSE, and MAE. An overview is provided in Table 2.

**Table 2:**
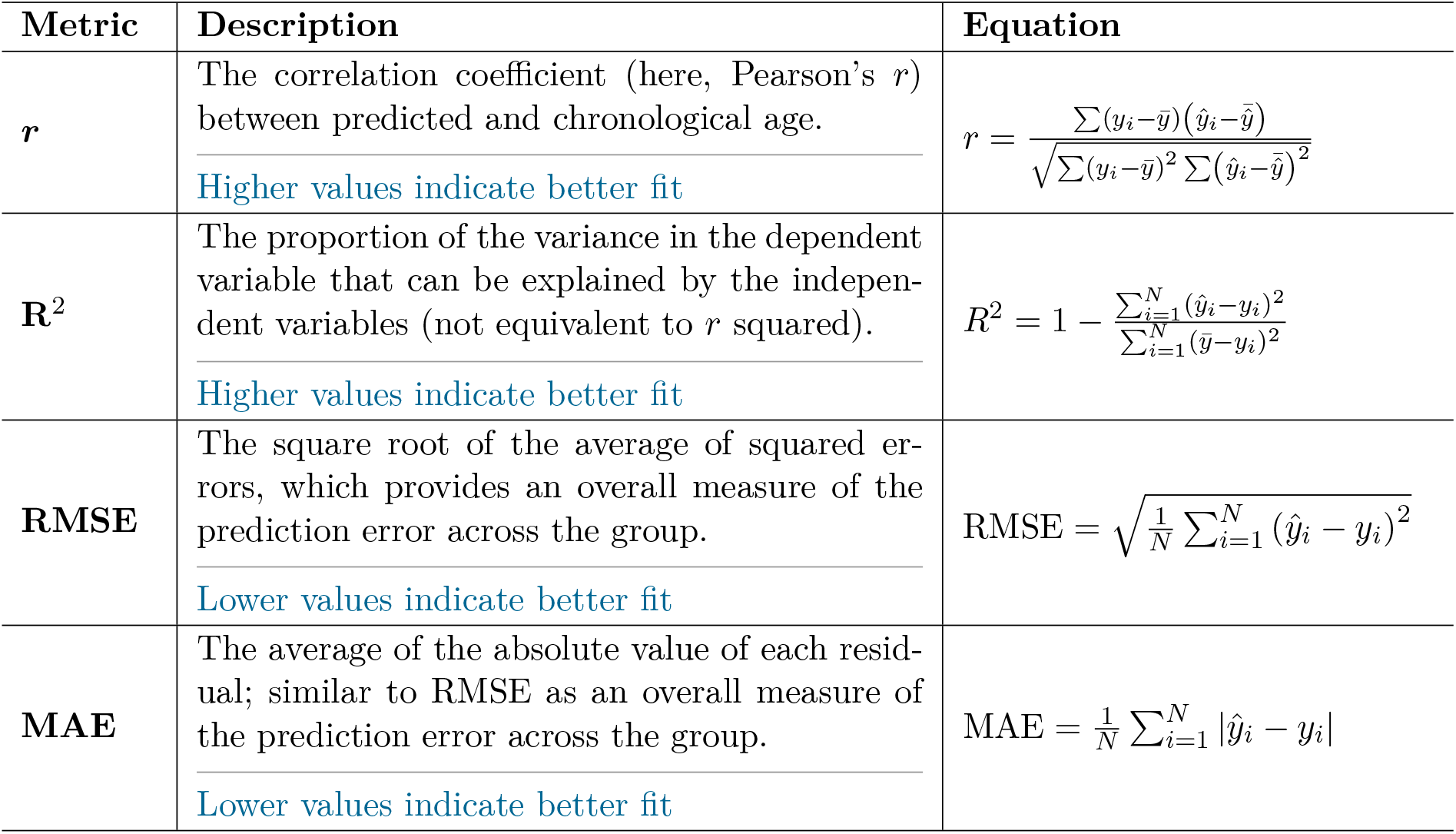
Overview of the model performance metrics and how they are usually interpreted in the context of model accuracy (blue font). RMSE = Root Mean Squared Error; MAE = Mean Absolute Error. Here, *y* are the true age values for each subject, *ŷ* are their predicted age values, 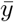 is the mean true age of the sample, and 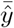 is the mean predicted age of the sample.

### 2.5. Effects of age range and sample size

To assess the effects of age range and sample size, we first ran 10-fold cross validated models within the full UKB and Cam-CAN samples to compare performance metrics between the two cohorts. Due to the large sample size, UKB data were used to systematically assess effects of age range in the subsets described below. Unless otherwise stated, sample size was held constant across training and test sets with N representing the maximum number of participants available with the narrowest age range.

#### 2.5.1. Test sets with varying age ranges, training set held constant

To assess the performance metrics in test sets with different age ranges, we trained a model on a subset including the full age range, and applied it to unseen test sets with different age ranges. In this setting, age range varies only for the test sets.

#### 2.5.2. Training sets with varying age ranges, test set held constant

To assess the performance metrics when age range was varied only for the training sets, we trained models based on subsets with different age ranges, and applied them to the same test set.

#### 2.5.3. Training and test sets with equal age ranges

To assess the performance metrics when age range was equal for training and test sets, we ran models using 10-fold cross validation within a series of subsets with different age ranges. To test the effects of age range in addition to sample size, we re-ran the 10-fold cross validation models using fractions of 2.5, 5, 10, 25, 50, 75, and 100 percent of the maximum number of participants available with the narrowest age range.

### 2.6. Age-bias correction

Brain-predicted age is often overestimated in younger subjects and underestimated in older subjects due to general statistical features of the regression analysis [38]. This phenomenon can be explained by the limiting case where a model is unable to predict age based on the input features. In this scenario, all subjects will be predicted to have the median age (equivalent to the mean age if the data are symmetrically distributed), because such an estimate minimises the residuals; this is the aim of regression/ordinary least squares fitting. Assigning the median age as the prediction for all subjects will overestimate young subjects and underestimate older subjects (see Figure 9). With increasing prediction accuracy, the degree to which the model predicts median age is reduced, since the predictions move closer to true age. Hence, age-bias is less pronounced in models with high prediction accuracy, but will always be present to some extent since the relationship between brain characteristics and age is not perfect (as in *x* = *y*). To account for the method-inherent age-bias, a statistical correction can be applied to the age predictions or brain age delta estimates [11, 12, 39, 41, 38, 55, 40, 33, 39, 56, 13]. An example of a correction procedure is provided in Figure 1, where a correction is applied to the predictions by first fitting *Y* = *α ×* Ω + *β*, where Y is the modelled predicted age as a function of chronological age (Ω), and *α* and *β* represent the slope and intercept. The derived values of *α* and *β* are used to correct predicted age with *Corrected Predicted Age* = *Predicted Age* + [Ω *−* (*α ×* Ω + *β*)]. This correction is equivalent to removing the effect of chronological age from either predicted age or brain age delta (see e.g. [42, 40, 38]). In other words, it provides the same results as regressing out chronological age from brain age delta and using the residuals [41, 6, 9, 25, 12], or including chronological age as a covariate in regressions/correlations between brain age delta and other variables of interest [41, 42, 49, 31].

**Figure 1:**
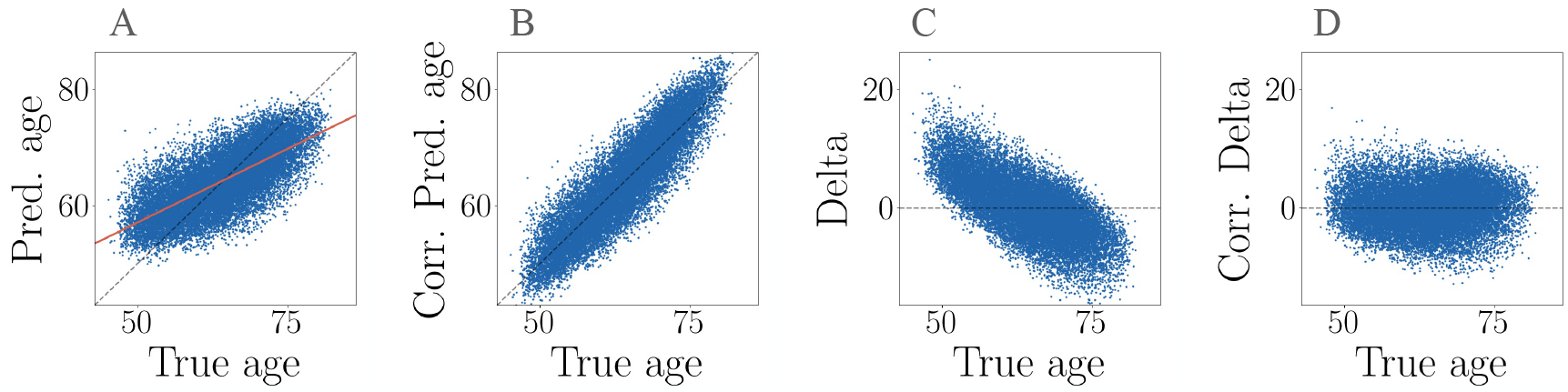
Example of age-bias correction. **A**: The uncorrected association between predicted and true age. The orange line shows the linear fit applied to model the age bias. **B**: The relationship between predicted and true age after using the coefficients from the fit (orange line in plot A) to correct predicted age. **C**: The uncorrected relationship between brain age delta and true age, illustrating the age dependence of delta. The negative slope is due to an anti-correlation between true age on the x-axis and negative true age on the y-axis, which occurs since negative true age is part of delta (*predicted age* − *true age*) **D**: Corrected delta calculated as *corrected predicted* − *age true age*, which shows no age dependence. Corrected delta obtained via a correction of the predicted age values gives equivalent results to correcting the delta values themselves for age [42]. Hence, while the corrected delta values show no age dependence, this is due to the alignment of corrected predicted age and true age as a result of the correction.

To assess the effect of age-bias correction on performance metrics, we applied the correction described above to i) the full UKB and Cam-CAN models, ii) UKB models based on subsets with different age range and sample sizes, and iii) a series of UKB and Cam-CAN models where 0, 10, 25, 50, and 75% of the data was randomly shuffled (age values are randomly reordered across subjects), to systematically assess corrected metrics across models with different levels of initial prediction accuracy.

As an alternative to simply regressing out age from brain age delta or correcting the predictions as described above, the coefficients from a fit in a training set can be used to correct the predictions or brain age deltas in an independent test set [40, 33, 39, 38, 56, 13]. With this approach, the correction in the test set is based only on the information about the age-bias observed in the training set. To test if this approach yielded different results, we split the full UKB and Cam-CAN samples in half to provide separate training and test sets, and corrected the predictions in the test sets based on coefficients derived from the fits in the training sets. The same cross-check was performed for the UKB models in section 3.3.

## 3. Results

### 3.1. Full models

The performance metrics for the 10-fold cross validated models including the total sample size and full available age range for each dataset are provided in Table 3. Despite the smaller sample size (622 versus 41,285 in UKB), the Cam-CAN prediction showed larger *r* and *R*^2^ values. The Cam-CAN model also showed larger RMSE and MAE values due to its wider age range (18 - 87 versus 45 - 82 in UKB). Hence, the lower RMSE/MAE values in UKB compared to Cam-CAN are not due to better model performance, but rather reflect that predictions in samples with a narrower age range are closer to the mean age of the group, which results in lower errors/smaller brain age delta values as shown in Figure 2. All performance metrics improved for both models after age-bias correction, as shown in Table 3. When correcting age-bias using fit coefficients derived from a training set to correct the predictions in a separate test set, the results were highly comparable (Supplementary Information (SI) Table 1).

**Table 3:** The correlations (*r*) between predicted age and chronological age, R^2^, root mean square error (RMSE), and mean absolute error (MAE) for the age predictions including the total sample and full age range in each of the datasets. The uncertainties for each parameter are also indicated. The performance metrics are provided before and after age-bias correction (corr).

|  | UKB | UKB corr. | Cam-Can | Cam-Can corr. |
| --- | --- | --- | --- | --- |
| $r$ | $0.722 \pm 0.002$ | $0.900 \pm 0.001$ | $0.875 \pm 0.008$ | $0.927 \pm 0.005$ |
| $R^2$ | $0.520 \pm 0.003$ | $0.767 \pm 0.002$ | $0.763 \pm 0.014$ | $0.837 \pm 0.011$ |
| RMSE [years] | $5.219 \pm 0.017$ | $3.640 \pm 0.012$ | $8.946 \pm 0.247$ | $7.415 \pm 0.191$ |
| MAE [years] | $4.188 \pm 0.015$ | $2.933 \pm 0.011$ | $7.182 \pm 0.217$ | $5.924 \pm 0.171$ |

**Figure 2:**
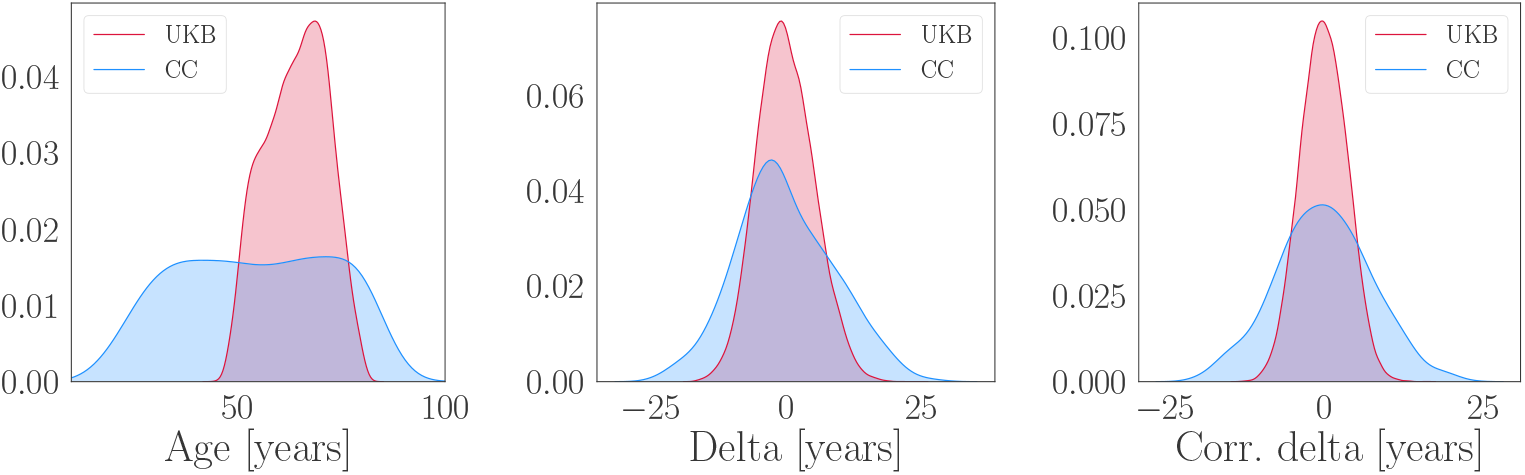
Age distributions (left plot), uncorrected brain age delta distributions (middle plot), and corrected brain age delta distributions (right plot) in UKB (red) and Cam-CAN (CC; blue). The distributions are normalised to have the same area, and the y-axes represent the density.

### 3.2. Effects of age range and sample size

This section shows model performance metrics measured in subsets with different age ranges. As a cross check, we repeated the age-range tests using samples where the lower instead of upper age limit was kept constant. The results were consistent, as shown in SI Figures 1-3.

#### 3.2.1. Test sets with varying age ranges, training set held constant

Figure 3 shows the model performance metrics calculated in UKB test sets with different age ranges when a model trained on the full age range is applied to each test set.

**Figure 3:**
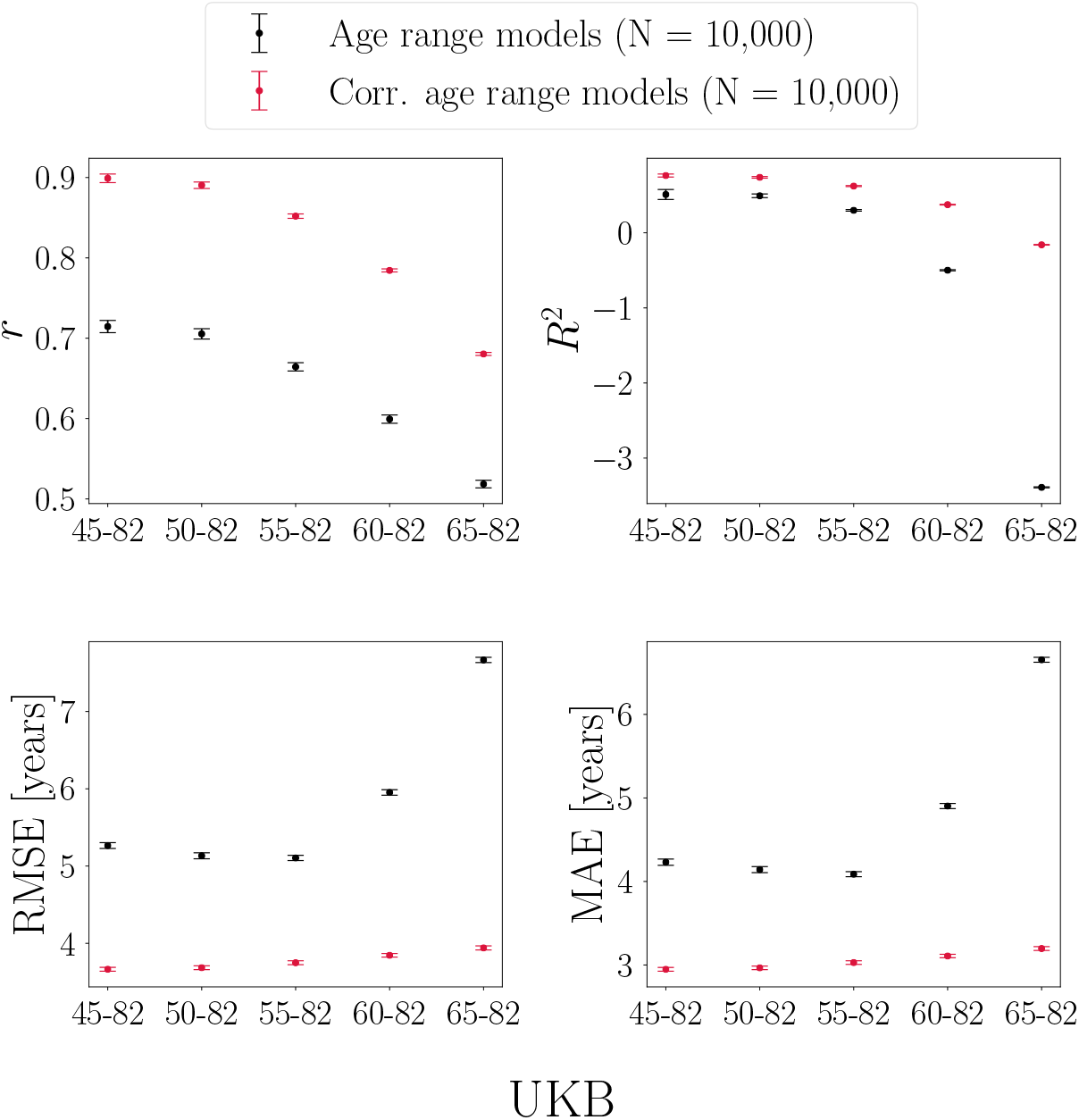
Performance metrics calculated in UK Biobank (UKB) test sets with different age ranges. Predictions are based on a model trained on the full age range. The x-axes indicate the age range for each of the test sets. Sample size is kept constant across training and test sets, and represents the maximum number of participants available with the narrowest age range (65-82y).

##### *r* and *R*^*2*^ values

As seen in Figure 3, *r* values are lower when calculated in test sets with a narrower age range, even though the predictions are based on a training set including the full age-range. The correlation coefficient is in general lower when measured in restricted ranges of a variable [36, 37], which is due to a smaller range in predicted and true age leading to less covariance. This also applies to the *R*^2^ values, but *R*^2^ is influenced by an additional effect; due to larger difference in mean age between the training and test sets, the *R*^2^ value becomes negative for the narrowest age range. The age-bias corrected *r* and *R*^2^ values are generally larger for all models, and the corrected values decrease with a narrower age range. In this scenario, the prediction variance is similar across test sets, which is a result of the training set being held constant. Hence, while both corrected and uncorrected *r* and *R*^2^ values are lower when measured in test sets with a restricted age range, low values do not imply that the brain-predicted age estimates are invalid (prediction variance is further discussed in Section 3.3). For *R*^2^, the test set with the narrowest age range shows the largest improvement after age-bias correction. This is because the correction adjusts the mean age difference between the training and test sets, as further described below.

##### RMSE and MAE values

As seen in Figure 3, RMSE and MAE initially decrease as the age range is narrowed, but then show a subsequent increase in the test sets with the narrowest age range. This trend is due to two competing effects: 1) the RMSE and MAE values generally *decrease* in test sets with a narrower age range due to smaller prediction range; 2) the RMSE and MAE values *increase* with a larger mean age difference between the training and test sets. When effect 2 becomes more prominent than effect 1, a turning point in RMSE and MAE is observed. The mean age and delta values for the training set and each of the test sets are shown in Table 4.

**Table 4:** Mean ± standard deviation for age and model errors/brain age delta values in the training set and each of the test sets with different age ranges. *Corr* indicates the age-corrected delta values. Larger mean age difference between training and test sets leads to smaller *R*^2^ values and larger RMSE and MAE values, as shown in Figure 3.

|  | Age | Brain age delta | Corr. Brain age delta |
| --- | --- | --- | --- |
| Training set (45-82y) | $64.08 \pm 7.52$ | $0.01 \pm 5.29$ | $-3.74 \times 10^{14} \pm 3.68$ |
| Test set (45-82y) | $64.17 \pm 7.52$ | $-0.08 \pm 5.26$ | $-1.85 \times 10^{14} \pm 3.66$ |
| Test set (50-82y) | $64.57 \pm 7.21$ | $-0.47 \pm 5.11$ | $-2.26 \times 10^{15} \pm 3.68$ |
| Test set (55-82y) | $66.22 \pm 6.10$ | $-2.09 \pm 4.66$ | $-3.37 \times 10^{17} \pm 3.75$ |
| Test set (60-82y) | $68.21 \pm 4.86$ | $-4.09 \pm 4.33$ | $6.47 \times 10^{15} \pm 3.85$ |
| Test set (65-82y) | $70.62 \pm 3.66$ | $-6.45 \pm 4.14$ | $1.05 \times 10^{14} \pm 3.94$ |

After age-bias correction, the RMSE and MAE values are generally smaller for all models, with similar values across test sets as seen in Figure 3. The similar values are due to stable prediction variance across test sets (a result of the training set being held constant). As seen for *R*^2^, the test set with the narrowest age range show the largest improvement in RMSE/MAE after age-bias correction, due to the adjustment of the mean difference between the training and test sets.

#### 3.2.2. Training sets with varying age ranges, test set held constant

Figure 4 shows the model performance metrics when models trained on different age ranges are applied to the same test set.

**Figure 4:**
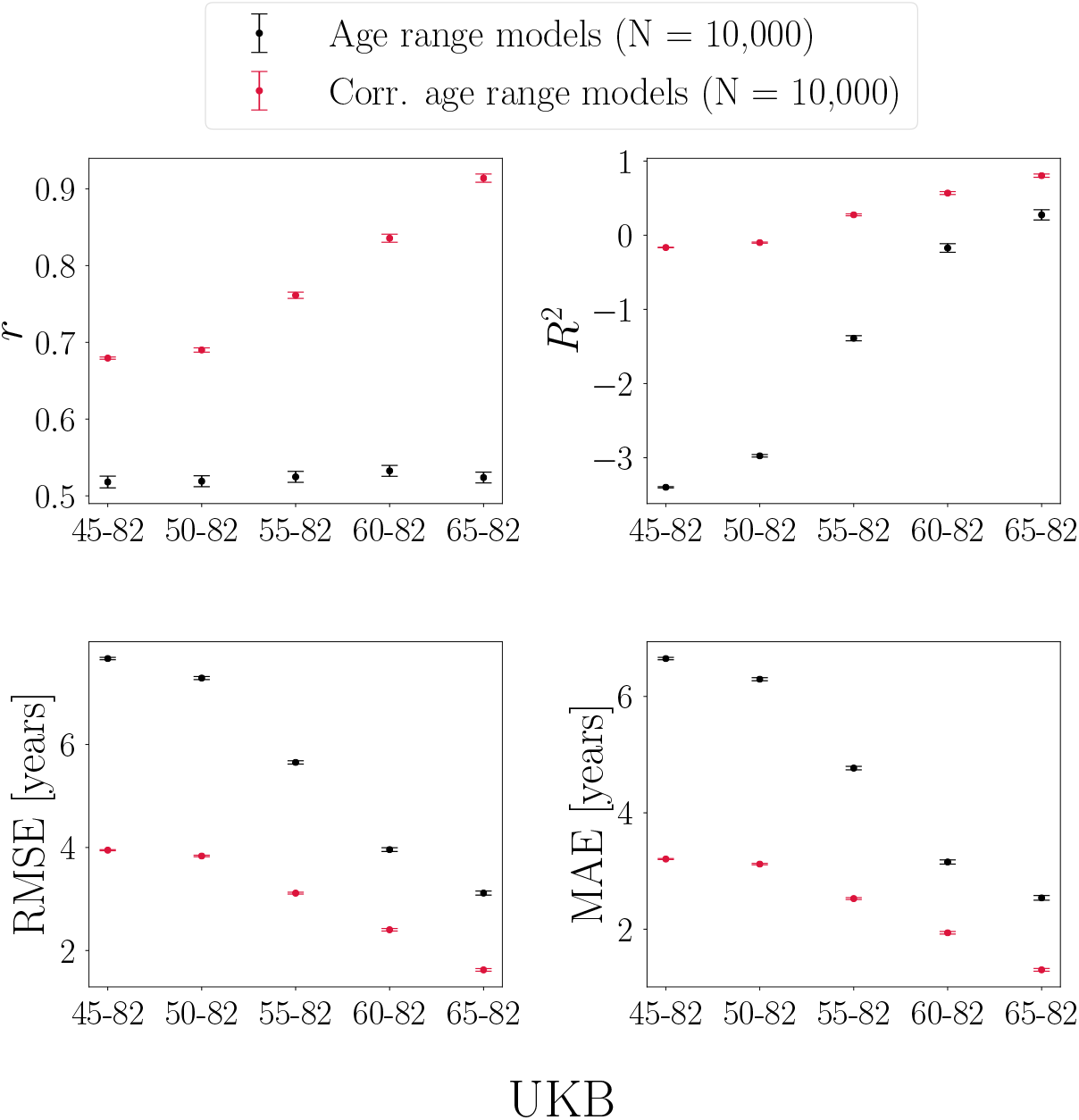
Performance metrics calculated in a UK Biobank (UKB) test set (age range = 65-82y). Predictions are based on models trained with different age ranges. The x-axes indicate the age range of the training sets applied to the same test set. Sample size is kept constant across training and test sets, and represents the maximum number of participants available with the narrowest age range (65-82y).

##### *r* and *R*^*2*^ values

As seen in Figure 4, the uncorrected *r* values are stable for all models, although the predictions are based on training sets with different age ranges. This is because the correlation coefficient is determined by the restricted age and prediction range in the test set (which is held constant). For *R*^2^, the uncorrected values increase substantially when the training is based on a narrower age range, due to the decreasing difference in mean age between the training and test sets (the mean age difference is largest when the training is based on the full age range, and smallest when the training is based on the narrowest age range (65-82y) as it matches the age range of the test set (65-82y)). After age-bias correction, the *r* values are generally larger for all models, but the largest improvement is seen for the model where the training is based on the narrowest age range. This is due to lower prediction variance in training sets with a narrower age range: the lower the initial variance, the larger the improvement in *r* after age-bias correction (see section 3.3). For *R*^2^, the largest improvement after age-bias correction is seen for the model where the training is based on the widest age range. This is because the correction adjusts the mean age difference between training and test sets, which is largest when the training is based on the widest age range.

##### RMSE and MAE values

As shown in Figure 4, RMSE and MAE decrease when the training is based on a narrower age range. This is due to two effects: i) lower prediction variance in models trained on a narrower age range, and ii) decreasing mean age difference between training and test sets. After age-bias correction, the largest improvements in RMSE and MAE are seen when the training is based on the widest age range. This is because the correction adjusts the difference in mean age between the training and test sets, which is largest when the training is based on the widest age range. Although the correction adjusts mean age differences, corrected RMSE and MAE values still decrease when training sets are based on a narrower age range. This is due to lower prediction variance with a narrower age range, which results in smaller model errors/brain age delta values (see Figure 8).

#### 3.2.3. Training and test sets with equal age ranges

Figure 5 shows the model performance metrics when 10-fold cross validations are run within different age-range subsets (age range is equal for training and test sets).

**Figure 5:**
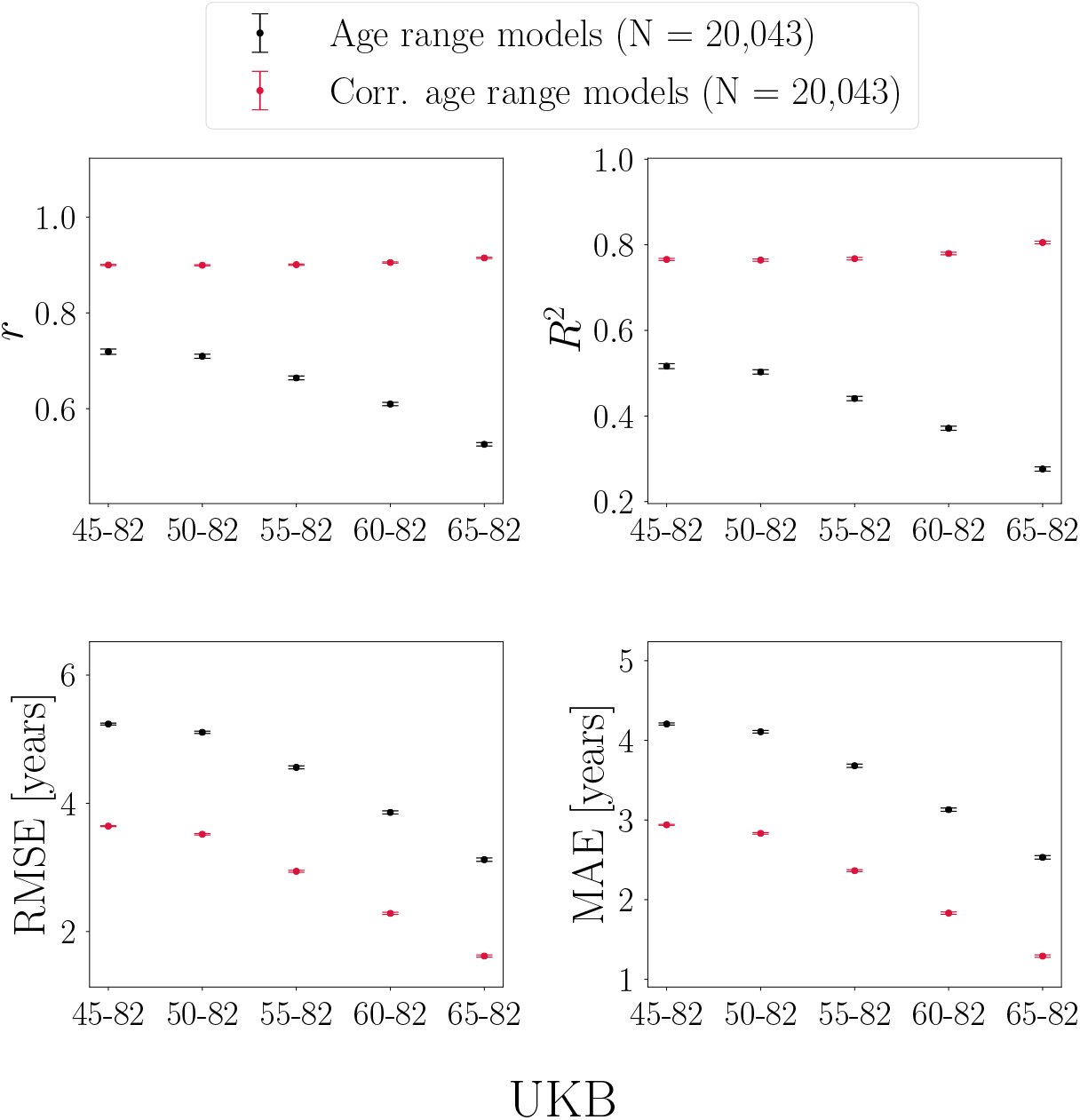
Performance metrics calculated in UK Biobank (UKB) subsets with different age ranges. Predictions are based on models trained using 10-fold cross validation within each subset, i.e. age range is equal for training and test sets. The x-axes indicate the age range for each of the subsets. Sample size is kept constant across subsets, and represents the maximum number of participants available with the narrowest age range (65-82y).

##### *r* and *R*^*2*^ values

As seen in Figure 5, the uncorrected *r* values decrease with a narrower age range. This is due to two effects: i) *r* is smaller in subsets with a narrower age range due to restricted age and prediction range, and ii) the variance in predictions is smaller when the training is based on a narrower age range. Since age range is equal for training and test sets within each subset, there are no mean age differences. Thus, *R*^2^ values are only influenced by the same effects as *r*; variable range and variance in predictions. After age-bias correction, the *r* values improve substantially across subsets, with the largest improvement seen for models with the lowest initial *r* values. This is due to lower prediction variance in subsets with a narrower age range (see section 3.3). The same effect is reflected in the corrected *R*^2^ values.

##### RMSE and MAE values

As shown in Figure 5, RMSE and MAE decrease with a narrower age range. This is due to the restricted prediction range in subsets with a narrower age range (predictions in samples with a narrower age range are closer to the mean age of the group, which equates to lower model errors/smaller brain age delta values). After age-bias correction, the RMSE and MAE values are generally smaller for all models, but the corrected values also decrease with a narrower age range. This is due to lower variance in subsets with a narrower age range, which results in smaller model errors/delta values (see Figure 8).

##### Effects of age range and sample size

As shown in Figure 6, all performance metrics improved with increasing sample size across subsets with different age ranges. Across all sample fractions, the effects of age range corresponded to the trends in Figure 5; lower uncorrected *r* and *R*^2^ values in subsets with a narrower age range due to restricted prediction range and lower variance, and lower RMSE and MAE values in subsets with a narrower age range due to restricted prediction range. Age-bias corrected metrics improved for all models, as shown in Figure 7.

**Figure 6:**
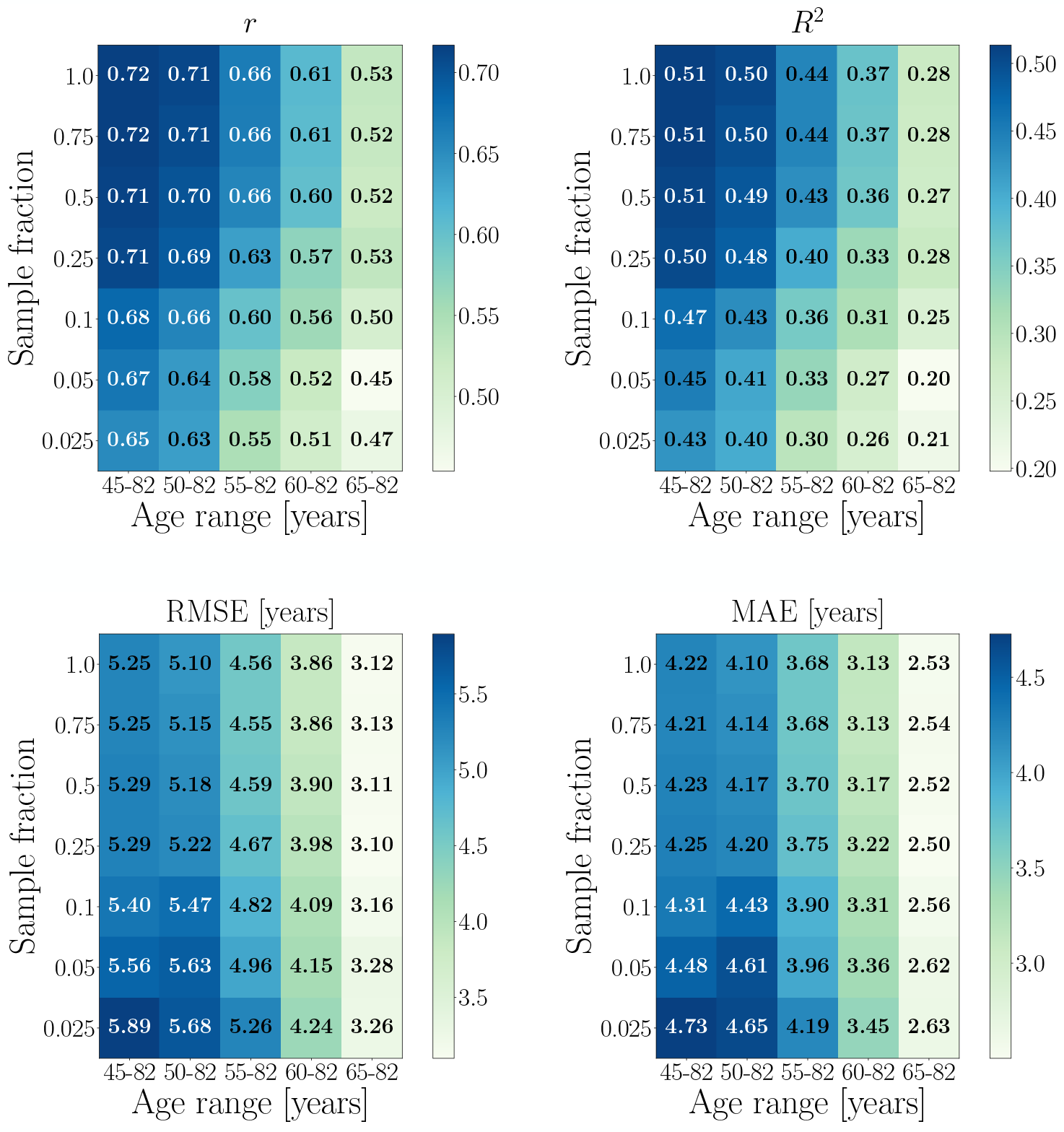
Performance metrics calculated in UK Biobank subsets with different age range and sample size. Predictions are based on 10-fold cross validation models run within each age-range subset, i.e. age range is equal for training and test sets within each subset. The x-axes show the age range for each subset, while the y-axes indicate the subset sizes in fractions of the maximum number of participants available with the narrowest age range; N for each sample fraction: 0.025 = 501, 0.05 = 1,002, 0.1 = 2,004, 0.25 = 5,011, 0.5 = 10,022, 0.75 = 15,032, 1 = 20,043.

**Figure 7:**
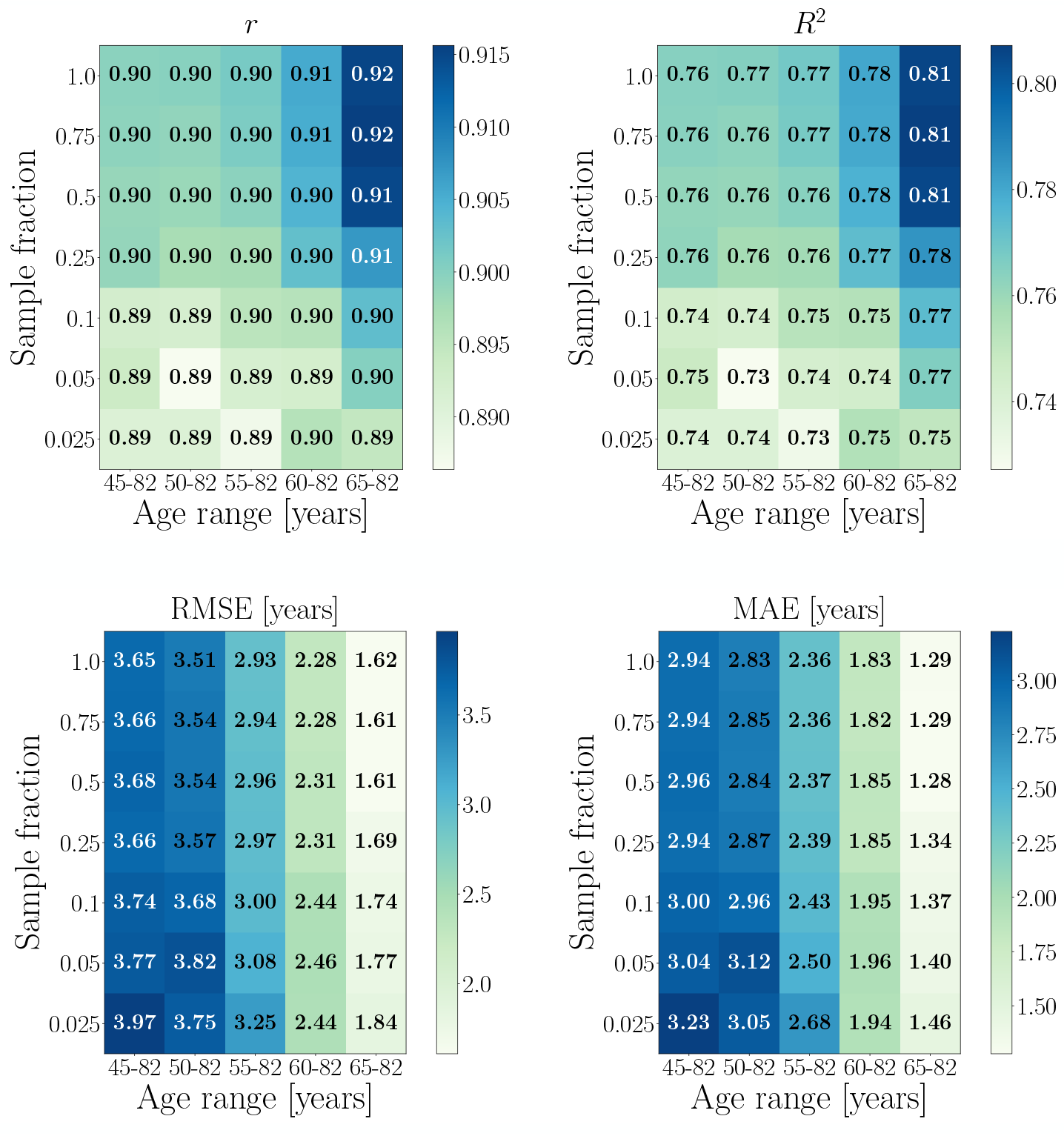
Age-bias corrected performance metrics calculated in UK Biobank subsets with different age range and sample size. Predictions are based on 10-fold cross validation models run within each age-range subset, i.e. age range is equal for training and test sets within each subset. The x-axes show the age range for each subset, while the y-axes indicate the subset sizes in fractions of the maximum number of participants available with the narrowest age range; N for each sample fraction: 0.025 = 501, 0.05 = 1,002, 0.1 = 2,004, 0.25 = 5,011, 0.5 = 10,022, 0.75 = 15,032, 1 = 20,043.

### 3.3. Age-bias correction applied to models with different levels of prediction accuracy

The results of applying the age-bias correction to models where 0, 10, 25, 50, and 75% of the data was randomly shuffled are shown in Figure 8. All performance metrics improved after correction, and the models with the poorest initial prediction accuracy (highest fraction of randomly shuffled data) showed the largest improvement after correction due to lower variance in predictions, as shown in Figure 9. The lower variance occurs with more predictions around the median age of the sample, which is a result of the model lacking sufficient information to provide accurate predictions. For Cam-CAN, all models improved to a similar extent after correction, as shown in SI Figure 4. The variance in the Cam-CAN data was more similar across models with different shuffle fractions (SI Figure 5) as compared to UKB, indicating that the wider age range provides more information for the model - leading to less predictions around median age.

**Figure 8:**
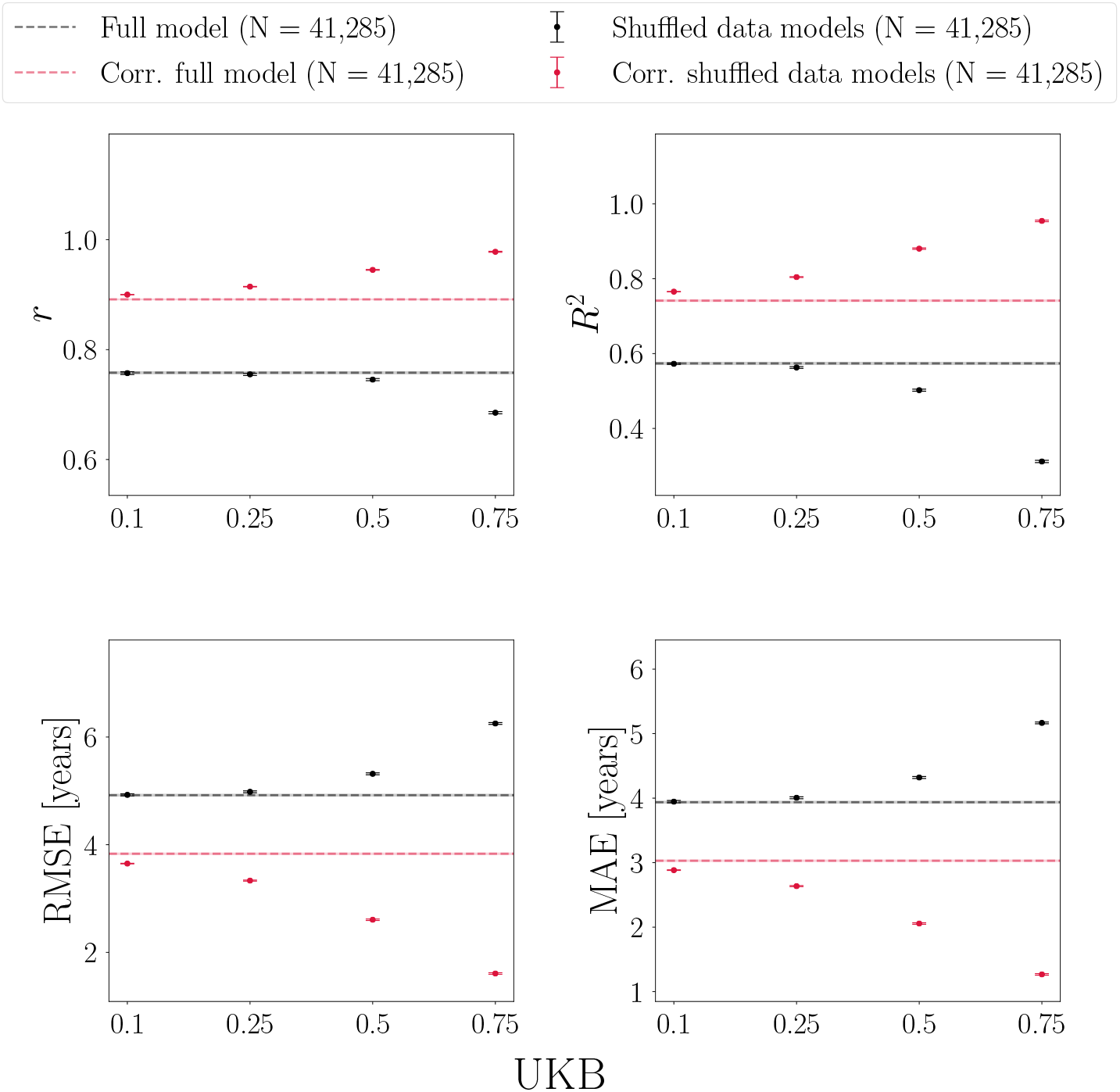
Age-bias correction in UK Biobank (UKB) models with 0, 10, 25, 50, and 75% randomly shuffled data. All models improve after correction, and the models with the poorest initial prediction accuracy (highest fraction of shuffled data) show the largest improvement. Hence, corrected metrics may not provide a relevant representation of initial model performance. Corr = corrected.

**Figure 9:**
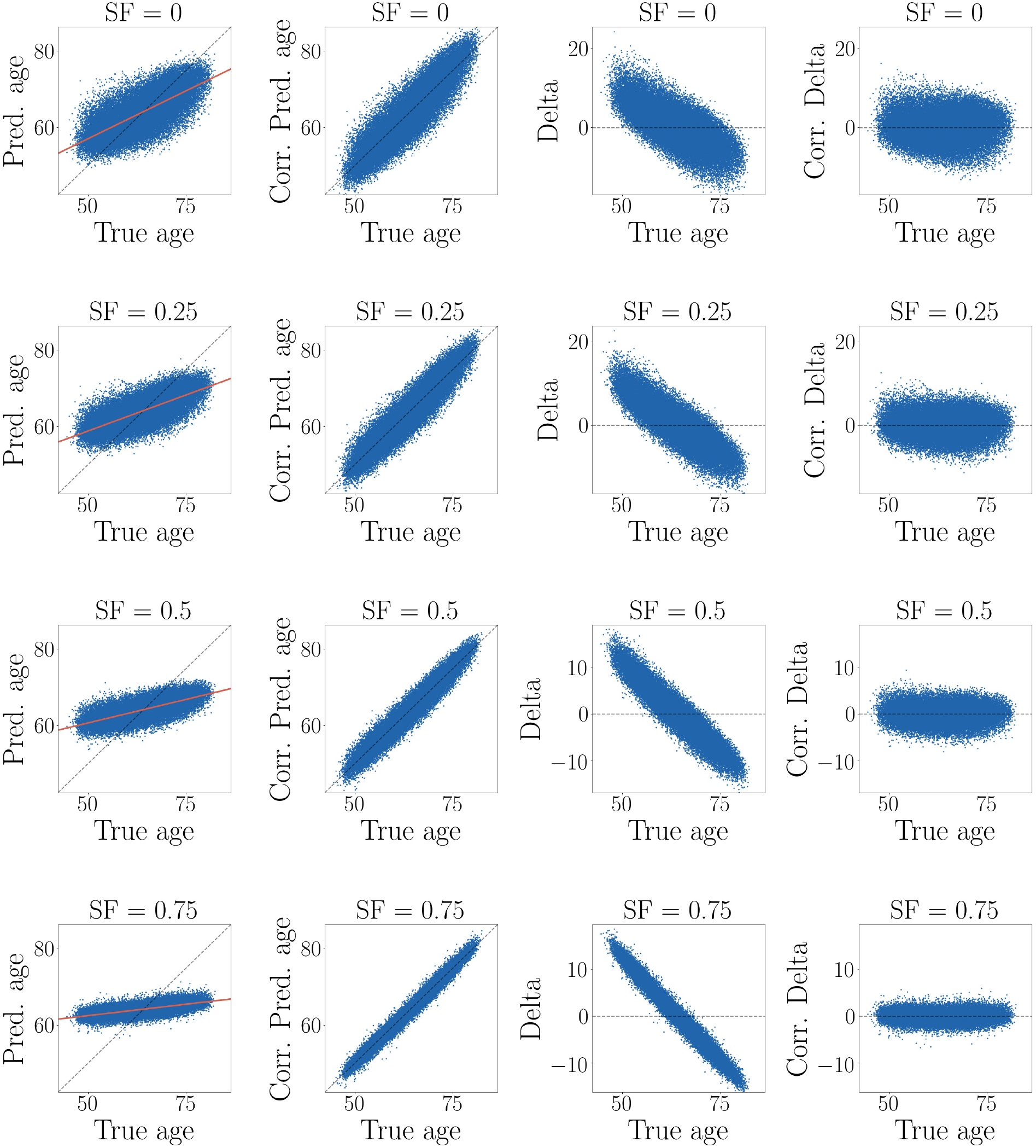
Age-bias correction in UKB models with randomly shuffled data. SF = shuffle fraction in %. **First column:** The plots of predicted versus true age show better performance for models with lower fractions of shuffled data. The models with the best performance also display the highest prediction variance, whereas the poorly performing models show predictions that cluster around median true age, resulting in low variance. **Second column**: The relationship between predicted and true age improves after age-bias correction, also for poorly performing models. **Third column:** Delta versus true age, illustrating the age dependence of delta. The negative slopes are due to an anti-correlation between true age on the x-axis and negative true age on the y-axis, which occurs since negative true age is part of delta (*predicted age*− *true age*). Models with smaller slopes in predicted versus true age (first column) show larger negative slopes in delta versus true age (third column) as a result of this. **Fourth column:** Corrected delta (*corr. pred age* − *true age*), which shows no dependence on age. Corrected delta obtained via a correction of predicted age gives equivalent results to correcting the delta values themselves for age [42]. Hence, while corrected delta shows no age dependence, this is due to a strong correlation between corrected predicted age and true age as a result of the correction (illustrated in SI Figure 15).

When using separate UKB training and test sets where the age correction parameters *α* and *β* were derived from a fit to the training set and used to correct the predictions in the test set, the results were highly comparable as shown in SI Figures 6 and 7. As a crosscheck, we repeated the age-bias analysis for UKB including a quadratic age term in the correction, which showed similar results (SI Figure 8).

### 3.4. UKB results based on SVR instead of XGB

The UKB results based on SVR instead of XGB are shown in SI section 6. In line with recent studies [38, 30], we found no evidence that choice of algorithm influenced the observed patterns: the effects of age range were highly comparable (SI Figures 9-11). The trends for subsets with different sample size and age range were also highly comparable, but XGB showed more stable performance across the smallest sample fractions (SI Figure 12). Age-bias correction showed equivalent effects for SVR and XGB models in samples where fractions of the data were randomly shuffled (SI Figures 13 and 14).

## 4. Discussion and summary of findings

Predicting age based on neuroimaging data can provide a useful marker for brain integrity and health [9, 24, 18, 3, 13], However, the current results emphasise that the model performance metrics *r, R*^2^, RMSE, and MAE cannot be directly compared across different studies, as they depend on factors including age range, sample size, prediction variance, and mean age differences between training and test sets.

### 4.1. Effects of age range

The results in section 3.2 show that model performance metrics depend on cohort age range in training and test sets. Since *r* and *R*^2^ values are lower when measured in restricted ranges of a variable [36, 37], these metrics can be lower when calculated in test sets with a narrow age range - also when the predictions are based on a training set with a wider age range. In this case, low *r* and *R*^2^ values are not indicative of poor model performance or insufficient variance in brain-predicted age estimates, but rather reflect the limited age variance in the test set. In studies where predictions are estimated in several sub-samples, it may be useful to include the age variance of the sub-sample with the largest age range in the calculation of performance metrics [19, 57], provided that the variances are similar in the sub-sample and a matching/restricted range of the sample used. In contrast, the use of training sets with a restricted age range can potentially involve poor model performance accompanied by low prediction variance, which is further discussed in Section 4.2.

In addition to age range and prediction variance, the *R*^2^ value is also influenced by differences in the mean age between training and test sets. Larger mean age differences lead to smaller *R*^2^ values, as well as larger RMSE and MAE values. However, the error metrics RMSE and MAE will in general *decrease* with a narrower age range, since predictions in samples with a narrower age range are closer to the mean age of the group (which results in lower model errors/smaller brain age delta values). Hence, small model errors do not necessarily reflect better model performance, and a model based on a cohort with a wide age range may show large *R*^2^ and *r* values accompanied by large RMSE and MAE values (as seen with Cam-CAN versus UKB in Section 3.1). Alternative model error metrics such as Relative Squared Error (RSE), Relative Absolute Error (RAE), Median Absolute Error, and weighted MAE also vary depending on age range, as shown in SI Section 8 (Figures 16-18).

### 4.2. Age-bias corrected versus initial model performance

The results in section 3.3 show how statistical age-bias corrections improve performance metrics by forcing an alignment between predicted and true age, leading to accurate predictions also for poorly performing models. This type of correction accounts for age-bias and mean age differences between training and test sets, but corrected performance metrics can also conceal potential issues with low prediction variance. While correcting the delta values instead of the predictions is common [40], these correction procedures lead to equivalent results [42] since the delta value contains the prediction minus age, and age is used in the correction fit (SI Figure 15). Hence, corrected deltas used in analyses to assess relationships with clinical or cognitive data are not exempt from the potential issues shown in Figures 8 and 9.

Inspection of uncorrected data can provide important information; for example, *r* and R^2^ values calculated in test sets with a narrow age range may be low, but prediction variance may be large if the training set has a wider age range. When the age range of the training set is also restricted, low *r* and R^2^ values may be due to low model performance accompanied by low prediction variance. Since age-bias corrected performance metrics do not contain information about these underlying model attributes, plotting the initial fit and data points can be helpful for evaluating the validity of brain-predicted age estimates. For example, if the relationship between the MRI input features and the dependent variable (age) is low in the training set, predictions may cluster around the median age of the sample as the model lacks sufficient information to provide accurate predictions. This would raise the question of what brain-predicted age estimates derived from models with low prediction accuracy actually represent, and whether other types of estimates (e.g. summary scores of the imaging data that are not obtained via age prediction) may be more appropriate in the given sample.

Since structural and functional brain measures show differential variation with age across the lifespan, age prediction accuracy varies depending on input features as well as cohort characteristics. For example, we found low age prediction accuracy based on resting-state functional MRI (fMRI) in UKB [58] and the Whitehall II MRI sub-study cohort (WHII) [10]. In WHII (N = 610, age range 60-85 years), the fMRI features showed weaker relationships with age compared to grey matter features derived from T1-weighted scans, and this result was also replicated in a matched UKB sub-sample in the same study. When systematically extending the UKB sub-sample, the fMRI prediction accuracy improved with a wider age range and larger sample size, but remained consistently lower than grey-matter based predictions in line with other UKB analyses [11]. Such findings further emphasise the challenges of comparing model results across studies, as model performance depends on specific brain characteristics and the age span over which they are modelled.

### 4.3. Conclusion

Performance metrics used for evaluating age prediction models depend on cohort and study-specific data characteristics, and cannot be directly compared across different studies. Although some effects can be mitigated through study designs where age distributions are carefully matched across training and test sets, observed model performance in a given test set cannot be generalised to samples with different age ranges. Since age-bias corrected metrics in general indicate high accuracy, even for poorly performing models, inspecting uncorrected results can provide important information about underlying model attributes such as prediction variance. While age prediction models have been used for more than a decade to generate imaging-based biomarkers [20], the approach continues to be developed and extended (see for example [3, 9, 56, 31, 49, 58]). Although not covered in the current study, an increasingly common scenario involves combining data from various cohorts and scanners, which poses additional challenges related to site- and scanner-dependent variance [25]. Improving methods for site/scanner adjustments [59, 60], or incorporating uncertainties into the predictions [61, 62], represent promising avenues for further developing robust and valid biomarkers for brain health and disease. As evident from the current results, clear reporting of sample characteristics and model attributes is important to enable accurate interpretation of model performance metrics in future work.

## Acknowledgements

This research was conducted using the UK Biobank under Application 27412. While working on this study, the authors received funding from the Swiss National Science Foundation (AM.G.dL.; PZ00P3_193658; B.D.; NCCR Synapsy, project grants Nr 32003B_135679, 32003B_159780, 324730_192755, and CRSK-3_190185), the Leenaards Foundation (B.D.), the Collaboratory on Research Definitions for Reserve and Resilience in Cognitive Aging and Dementia (M.A.; 5R24AG061421-03), the UK Medical Research Council (J.H.C. and M.A.; MR/R024790/2, K.P.E.; G1001354), the HDH Wills 1965 Charitable Trust (K.P.E.; 1117747), the research Council of Norway (L.T.W.; 273345, 249795, 223273; T.K.; 276082), the European Research Council under the European Union’s Horizon 2020 research and innovation programme (L.T.W.; 802998), the South-East Norway Regional Health Authority (L.T.W.; 2015073, 2019107), the German Research Foundation (K.F.; FR 3709/1-2; T.H.; HA7070/2-2, HA7070/3, HA7070/4), the Interdisciplinary Center for Clinical Research (IZKF) of the medical faculty of Münster (T.H.; MzH 3/020/20), the Interdisciplinary Center for Clinical Research (IZKF) of the Jena University hospital (K.F.; AMSP 07), and the ERA-Net Cofund through the ERA PerMed project “IMPLEMENT” (J.R.).

