## Supplementary Information (SI) Table 1 for "Mind the gap: performance metric evaluation in brain-age prediction"

### 1. Age-bias correction using separate training and test sets

SI Table 1: The correlations ($r$) between predicted age and chronological age, $R^2$, root mean square error (RMSE), and mean absolute error (MAE) for the age predictions in test sets including the full age range in each of the datasets. The uncertainties for each parameter are also indicated. The performance metrics are provided before and after age-bias correction (corr), where the predictions in the test sets are corrected based on the coefficients derived from a fit in the training set. N in training and test sets = 20,643 versus 20,624 for UKB, 311 versus 311 for Cam-CAN.

| | **UKB** | **UKB** corr. | **Cam-Can** | **Cam-Can** corr. |
|---|---|---|---|---|
| $r$ | $0.719 \pm 0.004$ | $0.900 \pm 0.001$ | $0.889 \pm 0.008$ | $0.930 \pm 0.005$ |
| $R^2$ | $0.516 \pm 0.005$ | $0.759 \pm 0.003$ | $0.790 \pm 0.014$ | $0.844 \pm 0.011$ |
| RMSE [years] | $5.215 \pm 0.026$ | $3.682 \pm 0.016$ | $8.427 \pm 0.234$ | $7.260 \pm 0.194$ |
| MAE [years] | $4.175 \pm 0.023$ | $2.971 \pm 0.015$ | $6.797 \pm 0.203$ | $5.788 \pm 0.171$ |

### 2. Effects of age range; lower instead of upper age limit kept constant

#### *2.1. Test sets with varying age ranges, training set held constant*

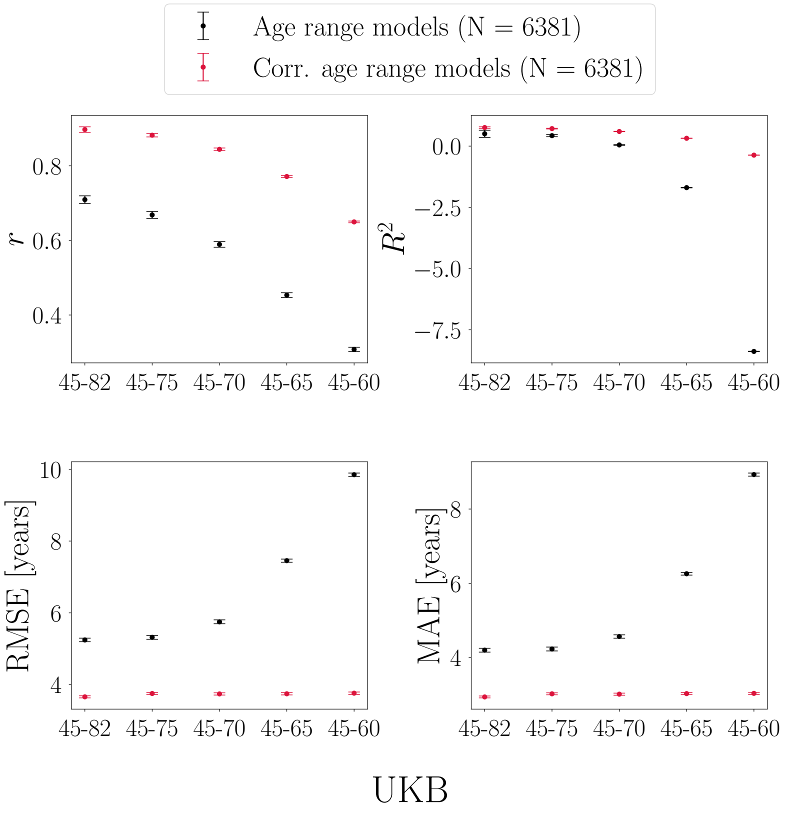


SI Figure 1: Performance metrics calculated in UK Biobank (UKB) test sets where the lower age limit was kept constant while the upper age limit was varied. Predictions are based on a model trained on the full age range. The x-axes indicate the age range for each of the test sets. Sample size is kept constant across training and test sets, and represents the maximum number of participants available with the narrowest age range (45-60y).

#### *2.1.1. Training sets with varying age ranges, test set held constant*

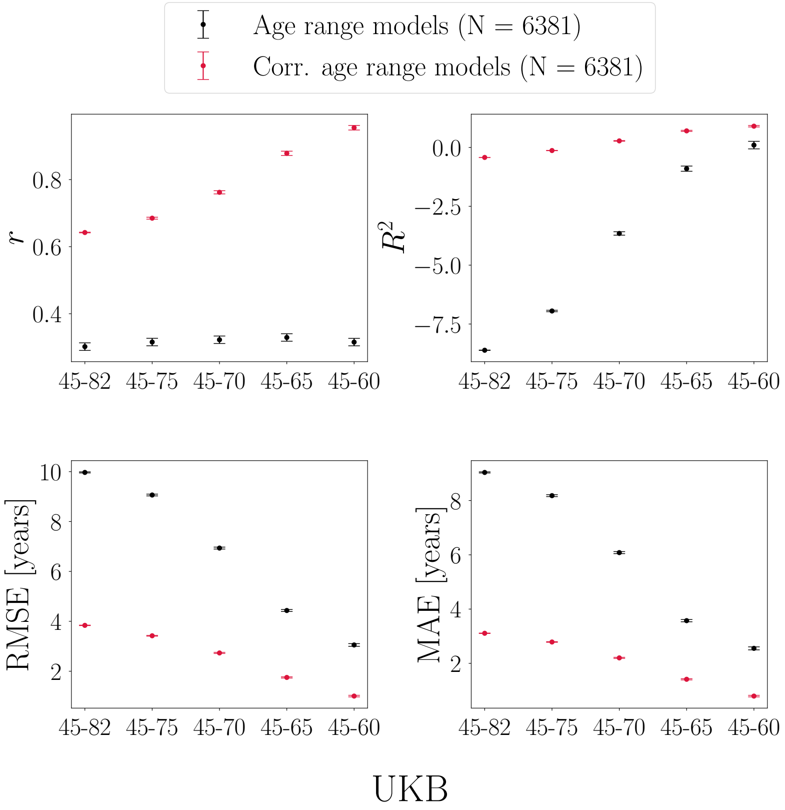


SI Figure 2: Performance metrics calculated in a UK Biobank (UKB) test set (age range = 65-82y). Predictions are based on training sets with different age ranges, where the lower age limit was kept constant while the upper age limit was varied. The x-axes indicate the age range of the training sets applied to the same test set. Sample size is kept constant across training and test sets, and represents the maximum number of participants available with the narrowest age range (45-60y).

#### 2.1.2. Training and test sets with equal age ranges

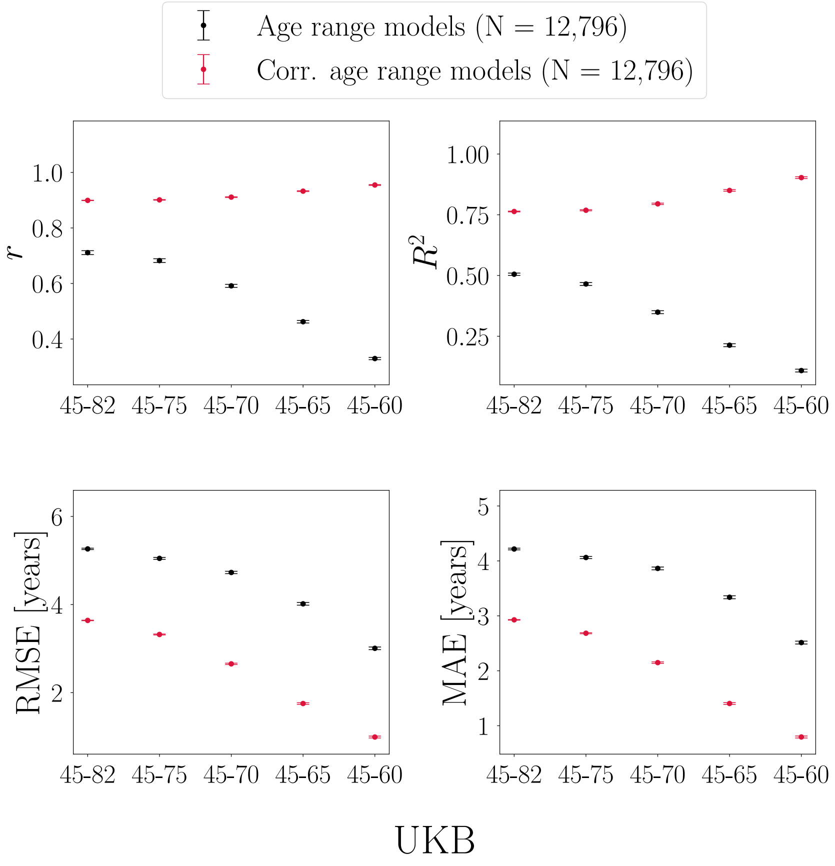


SI Figure 3: Performance metrics calculated in UK Biobank (UKB) subsets with different age ranges, where the lower age limit was kept constant while the upper age limit was varied. Predictions are based on models trained using 10-fold cross validation within each subset, i.e. age range is equal for training and test sets. The x-axes indicate the age range for each of the subsets. Sample size is kept constant across subsets, and represents the maximum number of participants available with the narrowest age range (45-60y).

### 3. Age-bias correction applied to models with different levels of prediction accuracy - Cam-CAN data

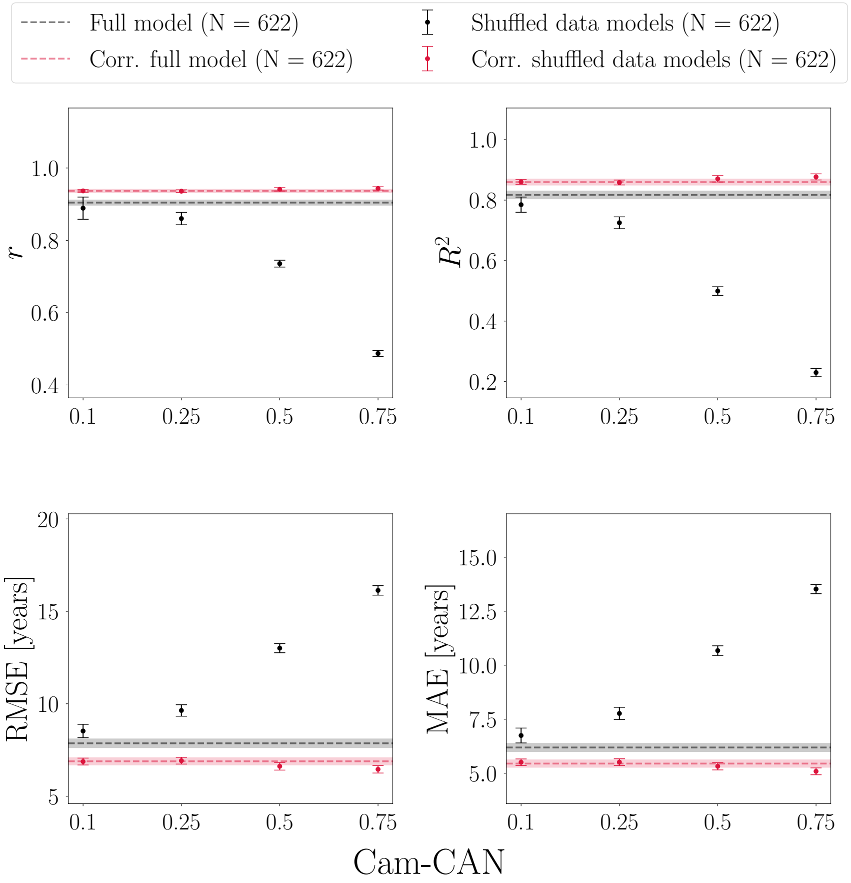


SI Figure 4: Age-bias correction in Cam-CAN models with 0, 10, 25, 50, and 75% of shuffled data. Corr = corrected.

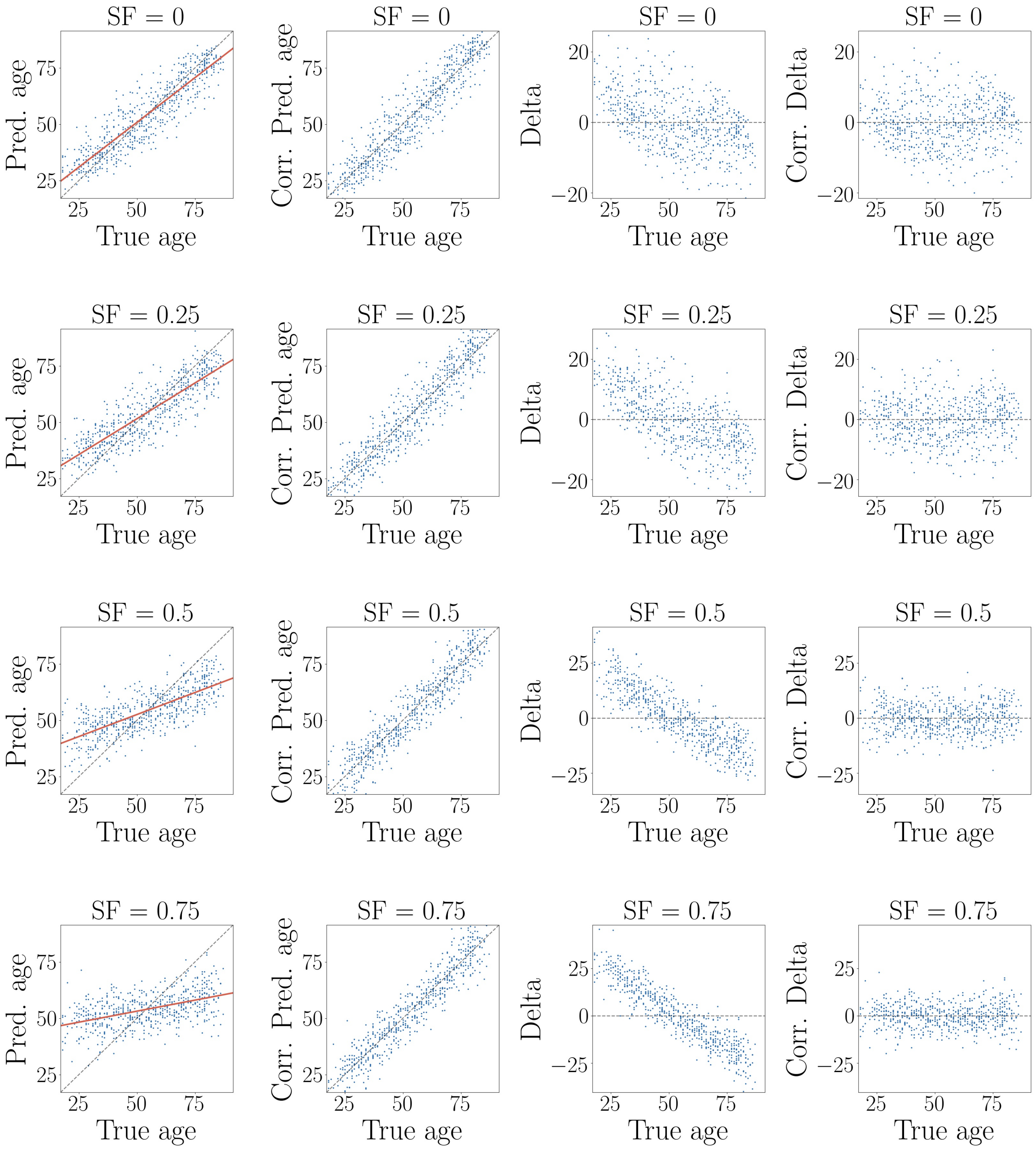


SI Figure 5: Age-bias correction in Cam-CAN models with 0, 25, 50, and 75% randomly shuffled data. SF = shuffle fraction in %. For all models, the relationship between predicted and true age improves after age-bias correction, and the delta values show a flat relationship with true age. Corr = corrected.

### 4. Effects of age-bias correction using separate training and test sets - UKB data

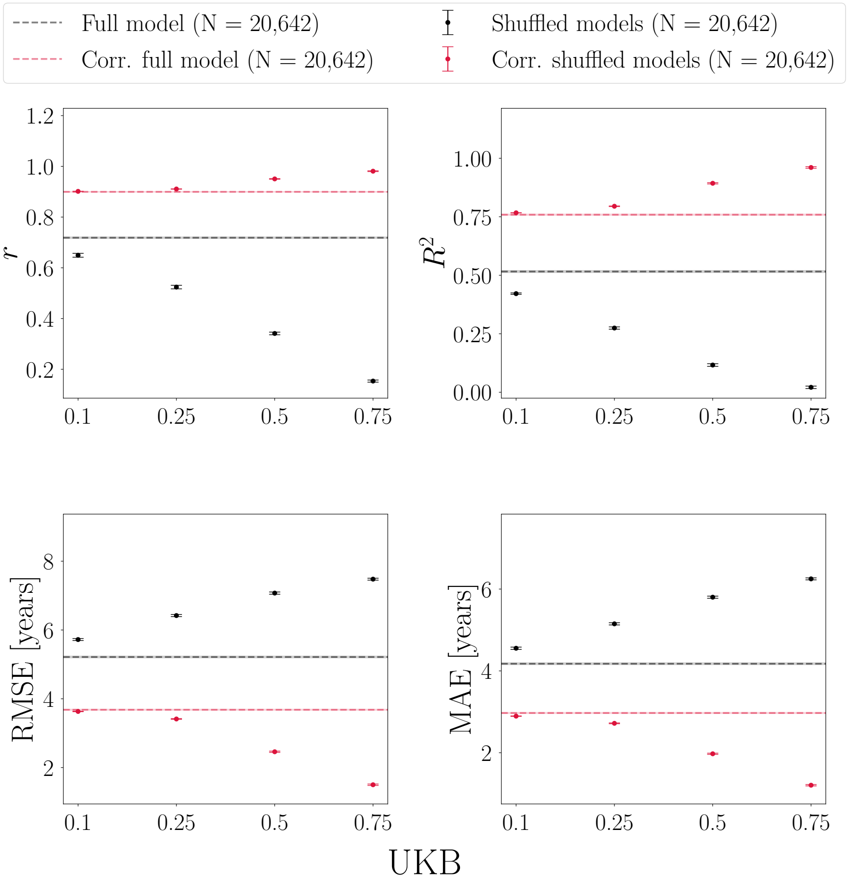


SI Figure 6: Age-bias correction with the correction fit applied to the predictions in a training set (N = 20,643), and the fit coefficients $\alpha$ and $\beta$ used to correct the predictions in a separate test set (N = 20,642) using UK Biobank (UKB) data. Performance metrics are shown for models with 0, 10, 25, 50, and 75% of shuffled data. All models improve after correction, and the models with the poorest initial prediction accuracy (highest fraction of shuffled data) show the largest improvement. Corr = corrected.

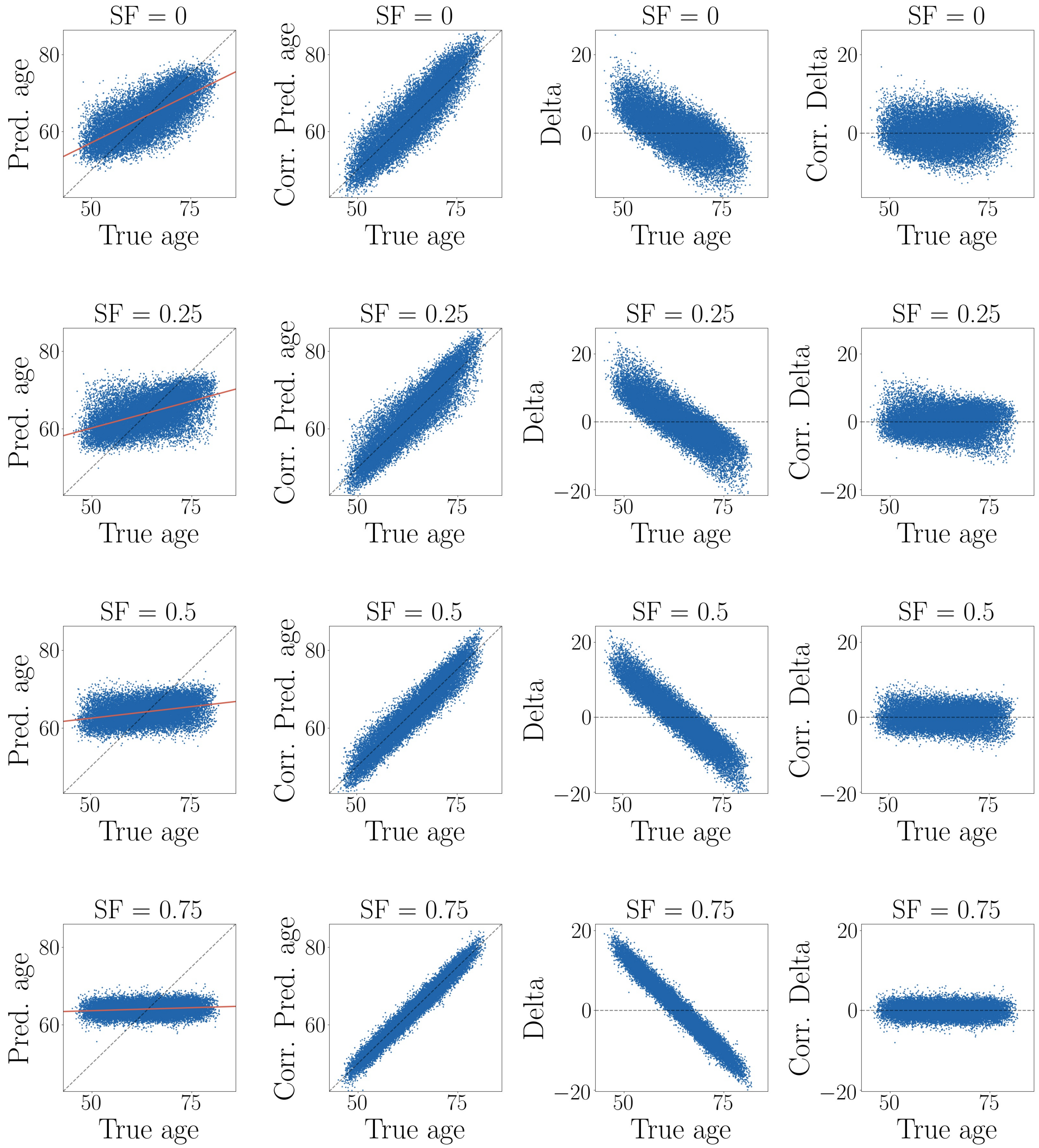


SI Figure 7: Age-bias correction with the correction fit applied to the predictions in a training set (N = 20,643), and the fit coefficients $\alpha$ and $\beta$ used to correct the predictions in a separate test set (N = 20,642) using UK Biobank data. Performance metrics are shown for models with 0, 25, 50, and 75% of shuffled data. For all models, the relationship between predicted and true age improves after age-bias correction, and the corrected delta values show a flat relationship with true age. The variance decreases with lower initial performance / higher shuffle fraction. Corr = corrected. For a detailed description of the plots, see Figure 9 in the main manuscript.

### 5. Effects of age-correction bias including a quadratic age term - UKB data

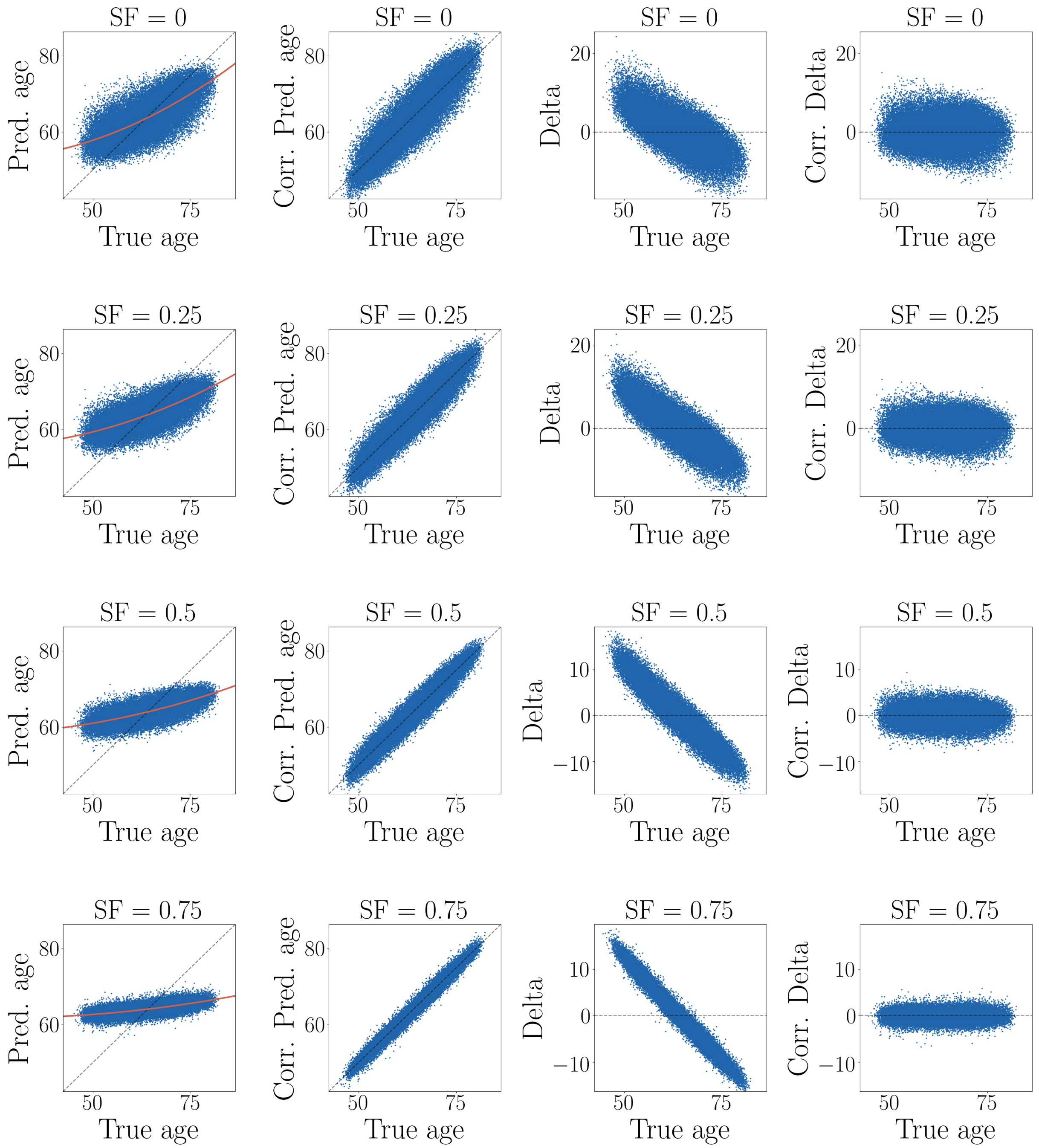


SI Figure 8: Age-bias correction including a non-linear term in UK Biobank models with 0, 25, 50, and 75% randomly shuffled data. SF = shuffle fraction in %. For all models, the relationship between predicted and true age improves after age-bias correction, and the corrected delta values show a flat relationship with true age. The variance decreases with lower initial performance / higher shuffle fraction. Corr = corrected. For a detailed description of the plots, see Figure 9 in the main manuscript.

### 6. UKB results using Support Vector Regression instead of XGBoost

#### *6.1. Test sets with varying age ranges, training set held constant*

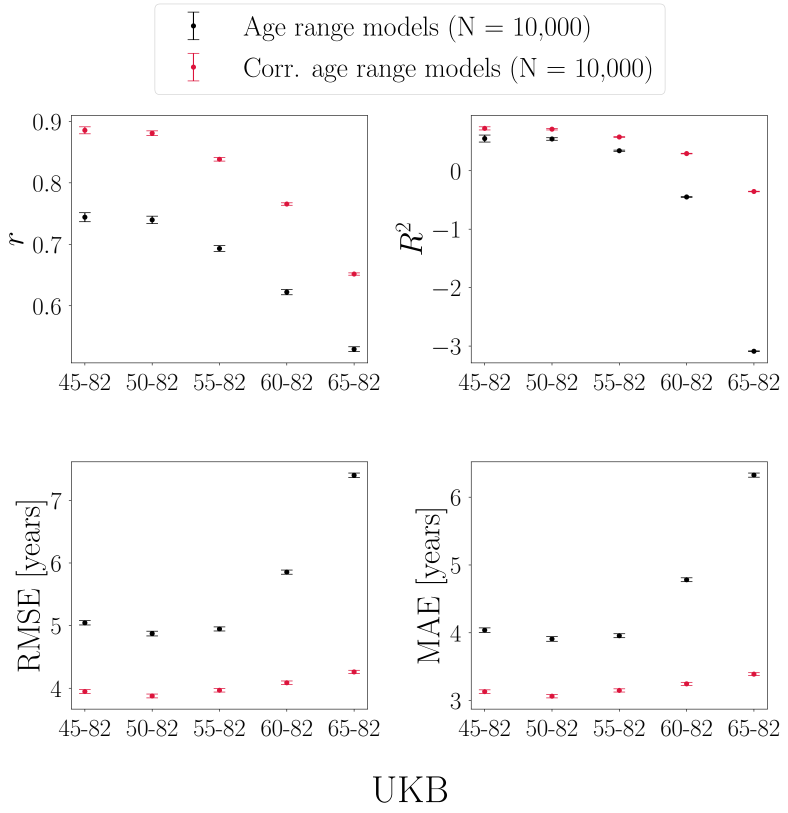


SI Figure 9: Performance metrics calculated in UK Biobank (UKB) test sets when using Support Vector Regression instead of XGBoost regression. Predictions are based on a model trained on the full age range. The x-axes indicate the age range for each of the test sets. Sample size is kept constant across training and test sets, and represents the maximum number of participants available with the narrowest age range (65-82y).

*6.1.1. Training sets with varying age ranges, test set held constant*

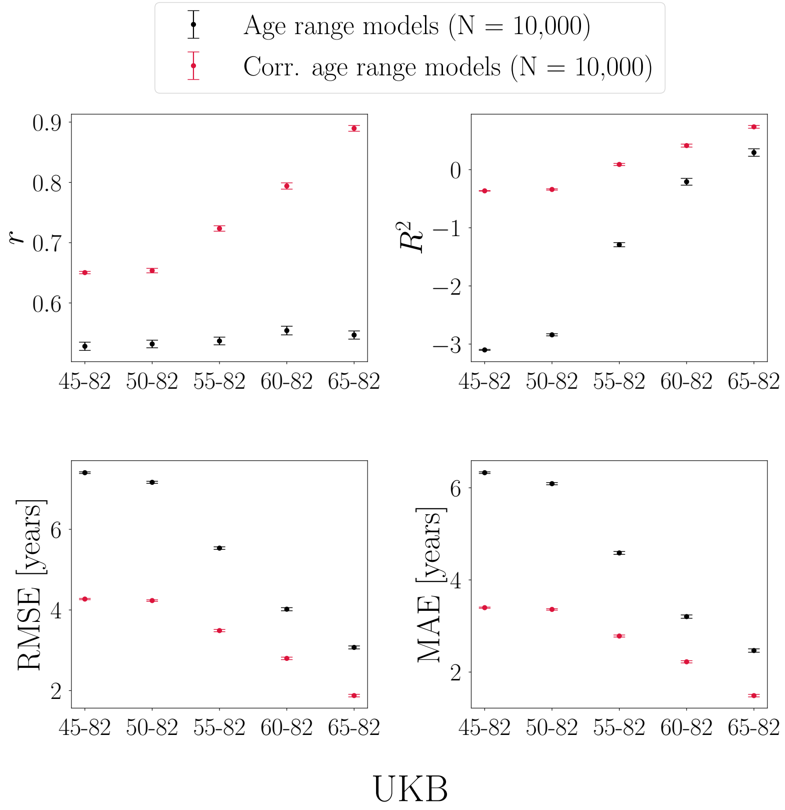


SI Figure 10: Performance metrics calculated in a UK Biobank (UKB) test set (age range = 65-82y) when using Support Vector Regression instead of XGBoost regression. Predictions are based on models trained with different age ranges. The x-axes indicate the age range of the training sets applied to the same test set. Sample size is kept constant across training and test sets, and represents the maximum number of participants available with the narrowest age range (65-82y).

#### *6.1.2. Training and test sets with equal age ranges*

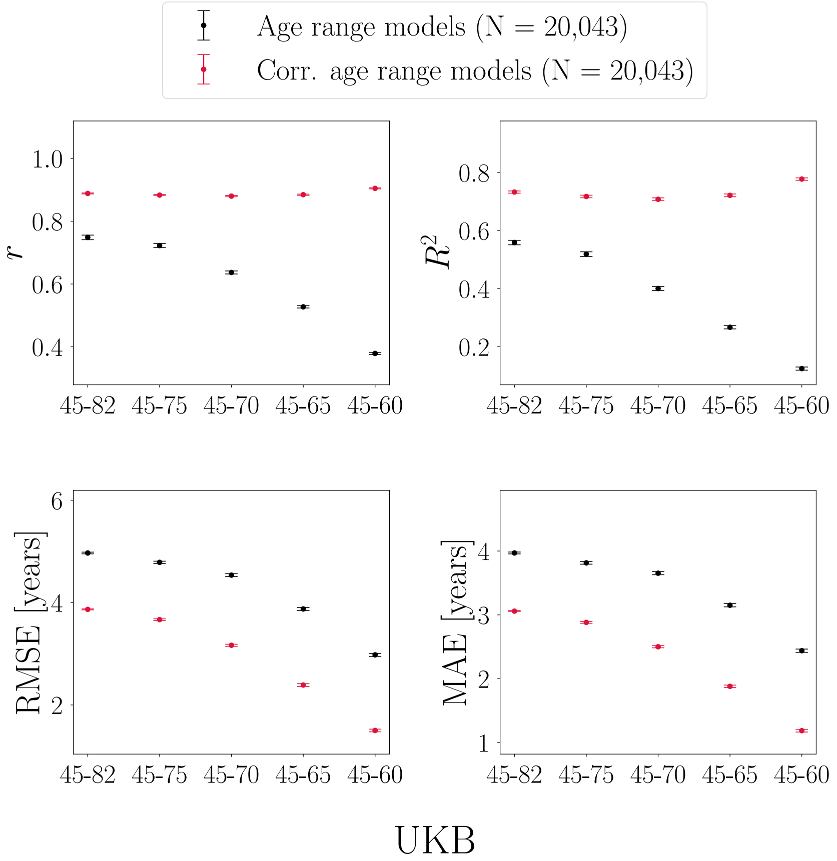


SI Figure 11: Performance metrics calculated in UK Biobank subsets with different age range and sample size when using Support Vector Regression instead of XGBoost regression. Predictions are based on models trained using 10-fold cross validation within each subset, i.e. age range is equal for training and test sets. The x-axes indicate the age range for each of the subsets. Sample size is kept constant across subsets, and represents the maximum number of participants available with the narrowest age range (65-82y).

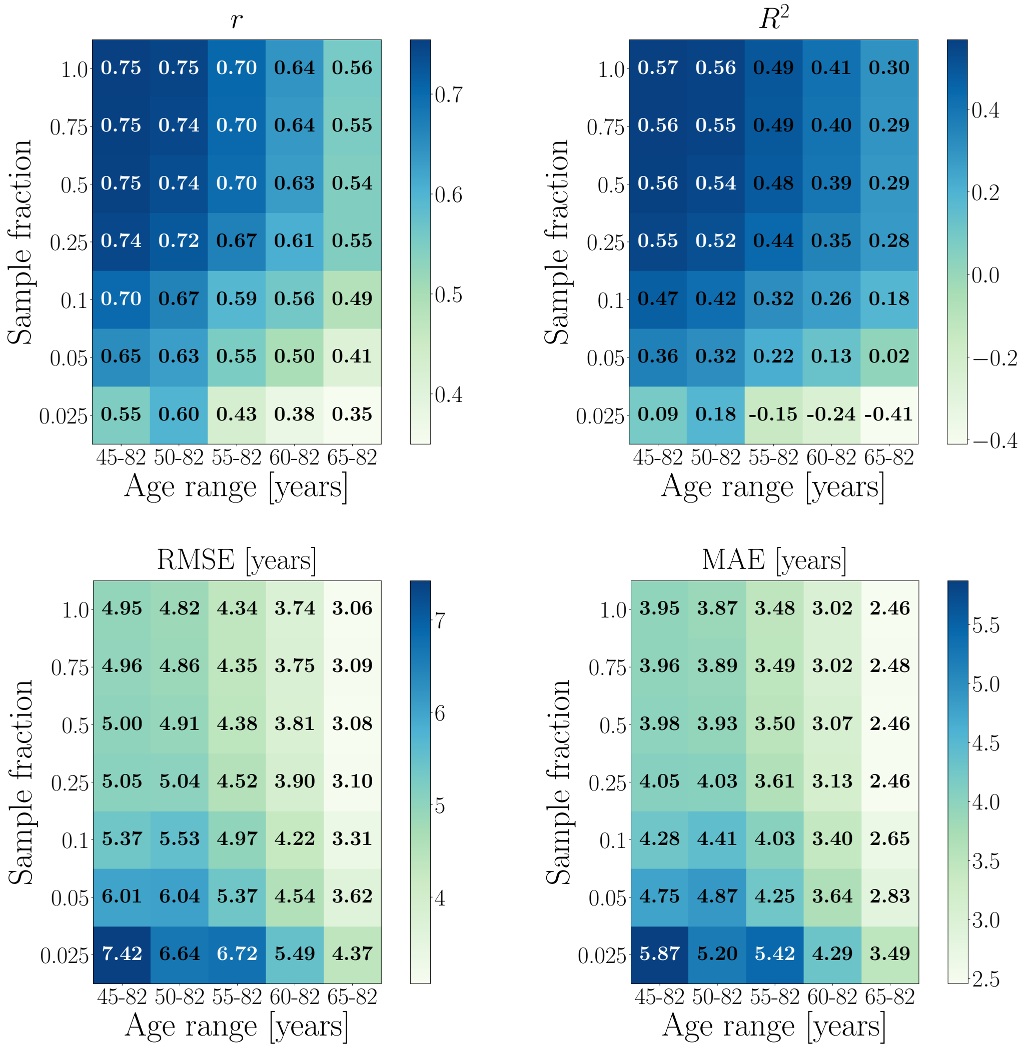


SI Figure 12: The effects of age range and sample size on model performance metrics when using Support Vector Regression instead of XGBoost regression. The x-axes show the age range for each subset, while the y-axes indicate the subset sizes in fractions of the maximum number of participants available with the narrowest age range; N for each sample fraction: 0.025 = 501, 0.05 = 1,002, 0.1 = 2,004, 0.25 = 5,011, 0.5 = 10,022, 0.75 = 15,032, 1 = 20,043.

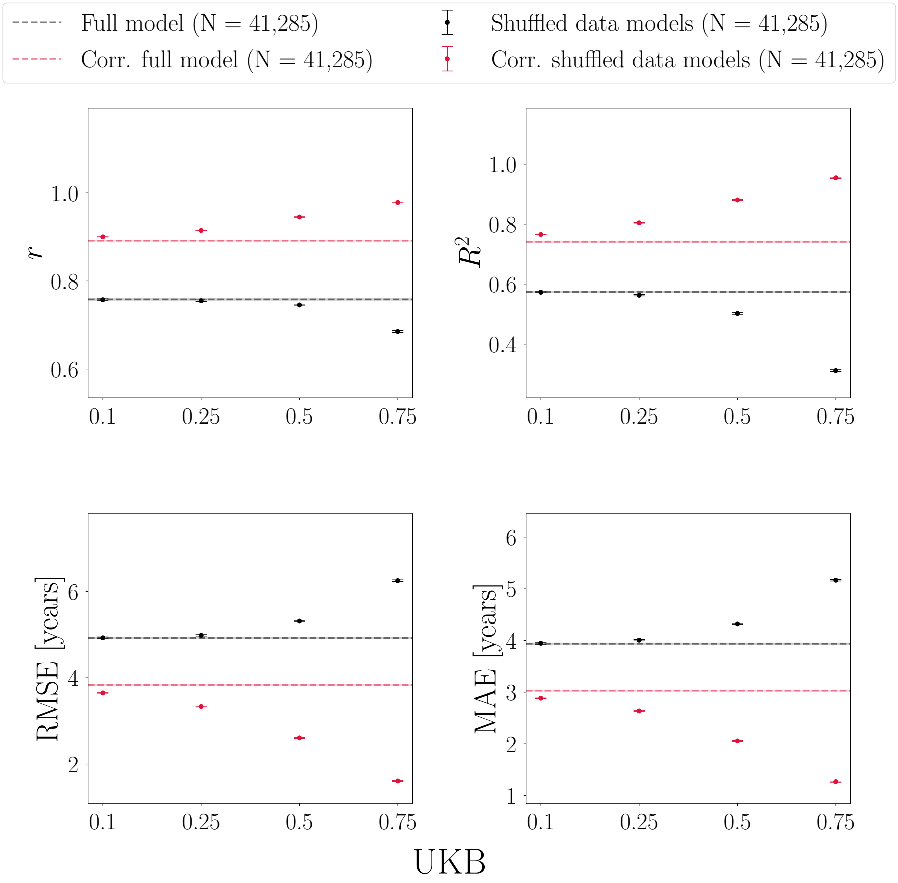


SI Figure 13: Age-bias correction in UK Biobank (UKB) models with 0, 10, 25, 50, and 75% randomly shuffled data, using Support Vector Regression instead of XGBoost regression. All models improve after correction, and the models with the poorest initial prediction accuracy (highest fraction of shuffled data) show the largest improvement. Corr = corrected.

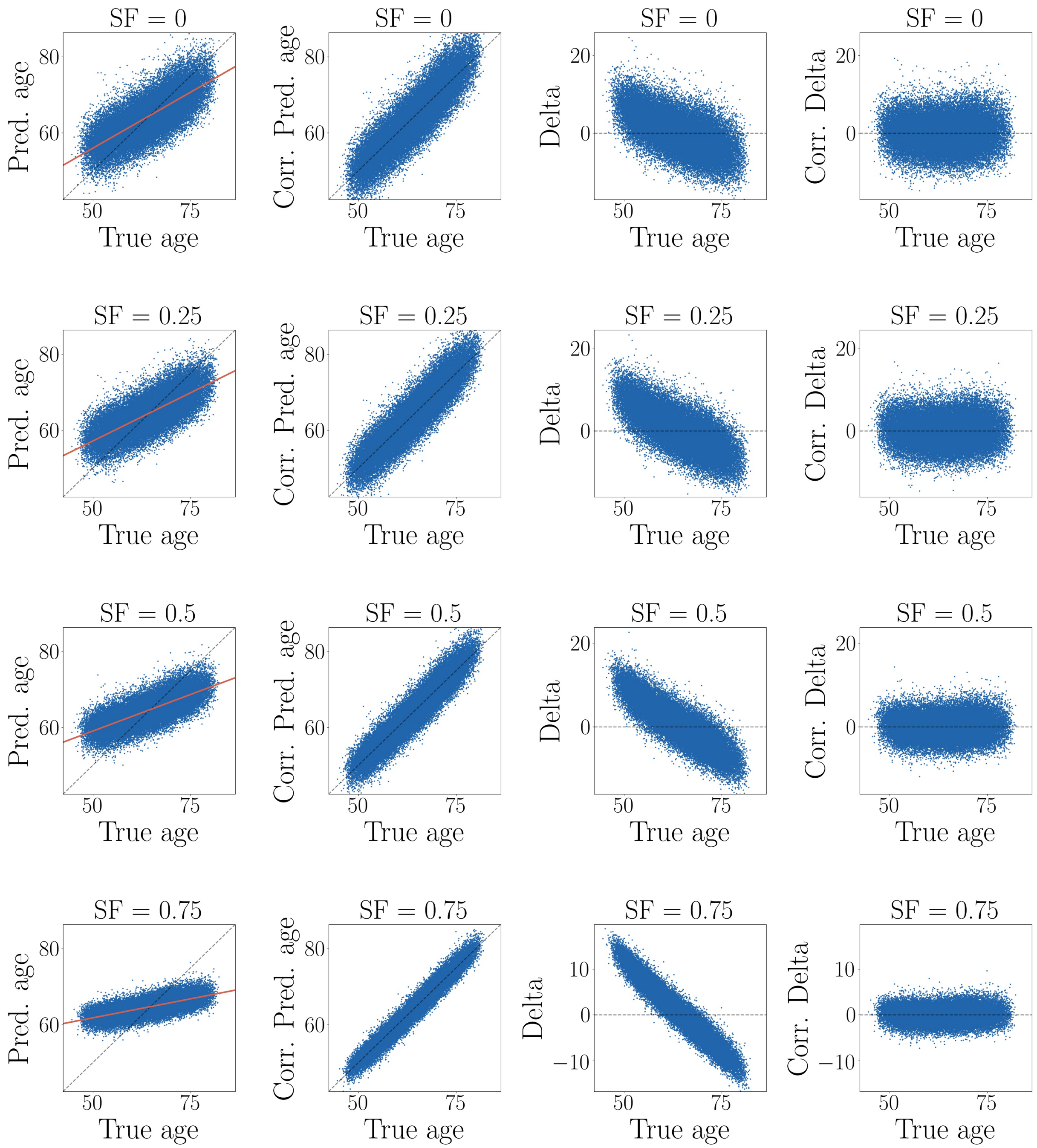


SI Figure 14: Age-bias correction in UKB models with randomly shuffled data when using Support Vector Regression instead of XGBoost regression. SF = shuffle fraction in %. For all models, the relationship between predicted and true age improves after age-bias correction, and the corrected delta values derived show a flat relationship with true age. The variance decreases with lower initial performance / higher shuffle fraction. Corr = corrected. For a detailed description of the plots, see Figure 9 in the main manuscript.

### 7. Age-bias correction applied to delta values instead of predictions - UKB data

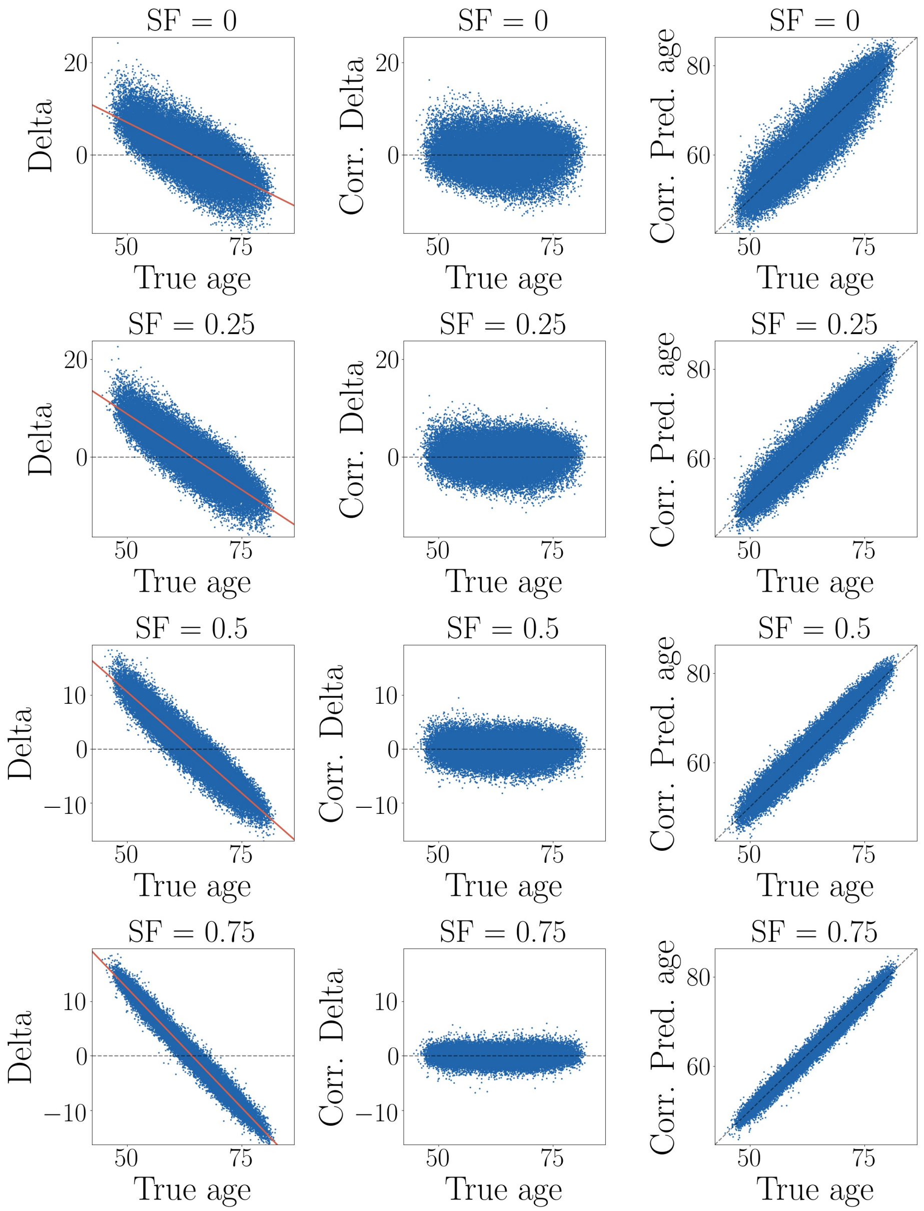


SI Figure 15: Age-bias correction applied to the brain age delta values instead of the predictions [1], shown for models with 0, 25, 50, and 75% randomly shuffled data in UKB. Delta is calculated as *predicted age* − *true age*. The deltas are corrected by fitting $Delta = \alpha \times \Omega + \beta$, where $\Omega$ represents chronological age, and $\alpha$ and $\beta$ represent the slope and intercept. If the corrected delta is subtracted from predicted age (third column), we see the strong relationships between predicted and true age as observed in Figure 8 in the main manuscript. Since the delta value contains the prediction minus age, and age is used in the correction fit, these correction procedures are mathematically equivalent [2], and corrected deltas are thus not exempt from the potential issues related to poorly performing models.

### 8. Alternative model error metrics

Alternative model error metrics such as Median Absolute Error (MedAE), weighted MAE (wMAE), Relative Squared Error (RSE), and Relative Absolute Error (RAE) also vary depending on age range, as shown in SI Figures 16-18. MedAE generally shows the same behaviour as MAE across different age ranges, but can be a useful metric as it is less affected by outliers. Weighted MAE is adjusted for the sample age range (calculated by MAE / age range [3, 4]), however, this metric only takes into account the range and not the underlying distribution. RSE is closely related to $R^2$, and can be expressed as RSE $= \sqrt{1 - R^2}$. For a model that describes a high proportion of the variance, RSE thus tends towards smaller values, whereas poorer models will have larger RSE values. RAE is related to MAE, but compares the prediction residuals $\hat{y} - y$ to the standard deviation of the predicted variable, $\bar{y} - y$. SI Figure 16 shows MedAE, wMAE, RSE, and RAE for a test sets with different age ranges based on a model trained on the full age range. As evident from the plots, these error metrics are influenced by variable range and mean age differences between training and test sets, similarly to RMSE and MAE.

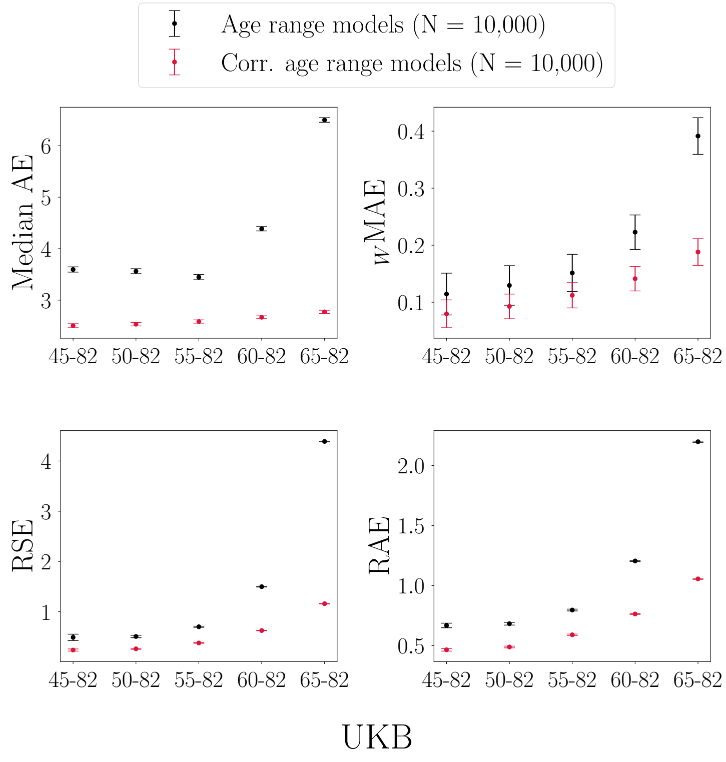


SI Figure 16: Alternative error metrics calculated in UK Biobank (UKB) test sets with different age ranges. Predictions are based on a model trained on the full age range. The x-axes indicate the age range for each of the test sets. Sample size is kept constant across training and test sets, and represents the maximum number of participants available with the narrowest age range (65-82y). wMAE = weighted MAE, RSE = Relative Squared Error, RAE = Relative Absolute Error.

SI Figure 17 shows MedAE, wMAE, RSE, and RAE calculated in a test set that is held constant while predictions are based on training sets with different age ranges. As evident from the plots, these error metrics are influenced by variable range, prediction variance, and mean age differences between training and test sets, similarly to RMSE and MAE.

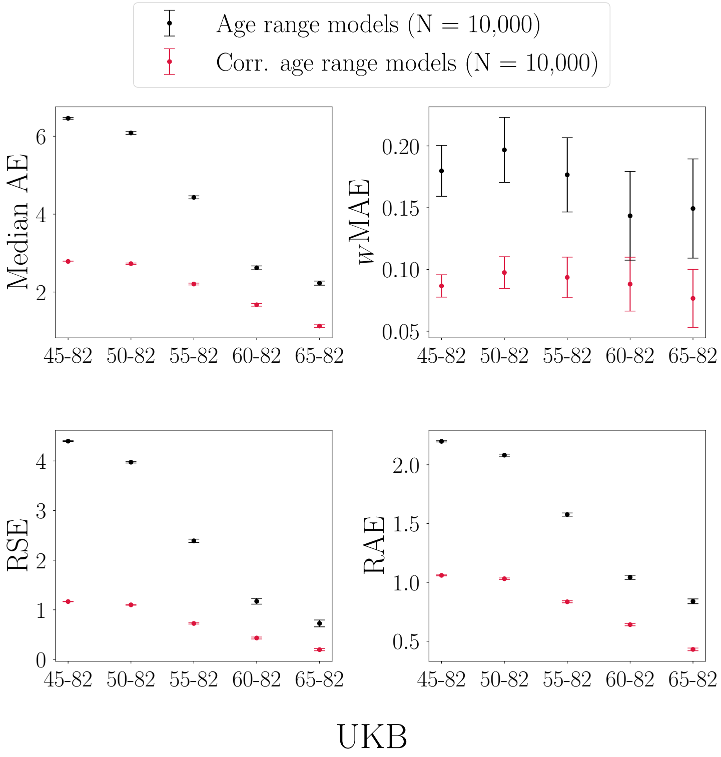


SI Figure 17: Alternative error metrics calculated in a UK Biobank (UKB) test set (age range = 65-82y). Predictions are based on models trained with different age ranges. The x-axes indicate the age range of the training sets applied to the same test set. Sample size is kept constant across training and test sets, and represents the maximum number of participants available with the narrowest age range (65-82y). wMAE = weighted MAE, RSE = Relative Squared Error, RAE = Relative Absolute Error.

SI Figure 18 shows MedAE, wMAE, RSE, and RAE calculated in subsets where 10-fold cross validations are run within different age-range subsets. In contrast to RMSE and MAE, the relative model error metrics wMAE, RSE, and RAE show *increasing* error values with a narrower age range, following the same trends as $R^2$ and $r$. This is due to how these metrics are calculated, where relative values are obtained by dividing by $\bar{y} - y$ (RSE and RAE) or the age range (wMAE). Here, there are no mean age difference between training and test sets, so the error metrics are influenced only by variable range and prediction variance.

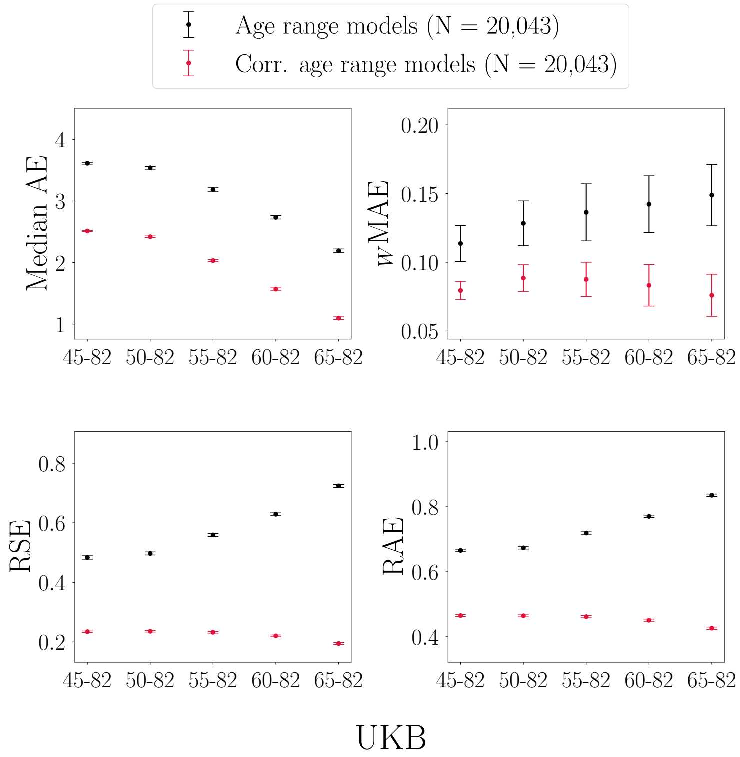


SI Figure 18: Alternative error metrics for models with different age ranges. Predictions are based on 10-fold cross validated models, i.e. age range is equal for training and test sets. The x-axes indicate the age range for each of the subsets. In contrast to RMSE and MAE, the error measures RSE, RAE, and weighted (w) MAE increase with a narrower age range (indicating larger model error), in line with the $r$ and $R^2$ patterns. This is due to how these metrics are calculated, where relative values are obtained by dividing by $\bar{y} - y$ (RSE and RAE) or the age range (wMAE). However, all metrics vary depending on the range of the predicted variable. wMAE = weighted MAE, RSE = Relative Squared Error, RAE = Relative Absolute Error.
